# Single-cell Characterization of DNA Hydroxymethylation of the Mouse Brain During Aging

**DOI:** 10.1101/2025.05.29.656780

**Authors:** Yali Bai, Tianjiao Yuan, Liuhao Ren, Yuhang Huan, Fa Yang, Yutong Li, Aicui Zhang, Yueyang Liu, Tian Tian, Ningxia Kang, Danni Chen, Xuchao Xie, Qiang Wang, Wei Chen, Yinghui Zheng, Xiaoliang Sunney Xie, Yunlong Cao

**Affiliations:** Changping Laboratory; Beijing, 102206, China; Biomedical Pioneering Innovation Center (BIOPIC), Peking University; Beijing, 100871, China; School of Life Sciences, Peking University; Beijing, 100871, China

## Abstract

DNA methylation dynamics, including 5-hydroxymethylcytosine (5hmC) and 5-methylcytosine (5mC), critically regulate brain function, yet conventional methods cannot distinguish these modifications, obscuring their distinct roles in gene regulation and aging. We present Joint- Cabernet, a bisulfite-free single-cell platform enabling simultaneous profiling of 5hmC, 5mC, and transcriptomes. Applying Joint-Cabernet to 84,071 nuclei from adult and aged mouse brains, we resolved cell-type-specific DNA hydroxymethylation landscapes, revealing elevated 5hmCG and 5hmCH levels in transcriptionally active genes across neuronal subtypes and spatial gradients in cortical layers. During aging, 5hmCG accumulates globally but is selectively enriched at open chromatin loci, aligning with the upregulation of cell-type-specific genes in distinct brain cell types. This single-cell DNA methylation brain cell atlas provides a framework for studying methylation-driven mechanisms in brain aging and neurodegenerative diseases.

## Introduction

The mammalian brain is a highly complex organ composed of diverse neuronal and glial cell types, each exhibiting specialized functions shaped by spatiotemporal gene expression programs (*1–4*). These programs are tightly regulated by epigenetic mechanisms, including DNA methylation, chromatin accessibility, and three-dimensional genome architecture (*5–10*). Among these, DNA 5-methylcytosine (5mC) and its oxidized derivative, 5-hydroxymethylcytosine (5hmC), play pivotal roles in neurodevelopment, synaptic plasticity, and aging (*11–16*). While 5mC is classically associated with transcriptional repression at heterochromatin and transposable elements, 5hmC accumulates in gene bodies of transcriptionally active neuronal genes, suggesting distinct functions for these modifications (*17–19*). However, the inability of conventional bisulfite-based methods to resolve 5mC and 5hmC has obscured their individual contributions to epigenetic dynamics, particularly in aging and neurodegenerative contexts.

Recent advances in single-cell methylome sequencing (e.g., snmC-seq3) and multi-omics profiling have begun to unravel cell-type-specific methylation landscapes (*6*). Yet scalable methods for simultaneous profiling of 5hmC, 5mC, and transcriptomes in individual cells remain lacking, limiting our understanding of how these marks coordinately influence gene expression and chromatin states across neuronal subtypes (*20*). Chemical and enzymatic approaches (e.g., TAB-seq, ACE-seq) have enabled base-resolution discrimination of 5hmC and 5mC in bulk tissues, but their application to single-cell analyses—especially in aging models—has been constrained by technical challenges in scalability and resolution (*21–27*).

Here, we present Joint-Cabernet, a high-throughput single-cell multi-omics platform based on our previous Cabernet method (*28*), that integrates transcriptomic, 5hmC, and 5mC profiling at single-cell resolution. Applying this method to 84,071 nuclei from adult and aged mouse brains, we resolved long-standing limitations of bisulfite-based methods and generated a DNA hydroxymethylation and methylation atlas of the mouse brain during aging, unveiling the distinct roles of 5hmC and 5mC in shaping brain function and aging-related DNA methylation remodeling.

## Results

### Technical development of Joint-Cabernet

Cabernet is an effective bisulfite-free method previously developed by our group for single-cell sequencing of 5mC and 5hmC, which involves the Tn5 transposome for DNA fragmentation followed by an enzymatic conversion protocol, including TET-mediated oxidation, BGT-mediated glycosylation, and APOBEC-mediated deamination (*28*). However, due to low uniformity, Cabernet’s direct Tn5 dual-barcoding using Tn5 transposome complexes preassembled with custom oligonucleotides prevents high-throughput single-cell profiling. To solve this problem, inspired by the s3 sequencing strategy (*29*), we implemented a template-switching-based barcoding strategy while retaining the Cabernet chemistry (fig. S1A). DNA fragmentation was performed with universal transposomes containing a cytosine-depleted reverse adapter, a uracil linker, and mosaic end (ME) sequences (table S1). During DNA fragmentation, uracil residues served as transient blocking sites during polymerase extension. Then, a secondary oligonucleotide directed template-guided extension to append barcodes and adapters while incorporating modified dCTP. These secondary oligonucleotides featured three functional domains: (i) a locked nucleic acid (LNA)-modified ME reverse complement (to elevate annealing specificity and suppress intramolecular self-looping), (ii) an inline cell barcode, and (iii) a forward adapter.

Thermal cycling was iterated over 10 rounds during extension to maximize processivity, ensuring near-complete conversion of fragmented templates into barcoded products. Critically, the forward adapter-bearing secondary oligonucleotide initiated extension toward the reverse adapter, yet uracil-induced termination sites strategically arrested elongation before completion. This blockade eliminated artifactual fragments that could spuriously incorporate exogenously methylated cytosines, thereby preserving the fidelity of endogenous methylation analysis. Under this scheme, the template-switching based extension produced fragments with a cytosine-depleted reverse adapter and cytosine-modified barcode and forward adapter, conferring resistance to APOBEC-mediated deamination.

The resulting 96 uniquely barcoded fragments were pooled prior to enzymatic processing steps following the Cabernet workflow (*28*). Specifically, one channel preserved both modifications using TET2 (to oxidize 5mC to 5-carboxylcytosine) and β-glucosyltransferase (BGT, to glycosylate 5hmC), while the other channel employed BGT alone to protect 5hmC. APOBEC3A then deaminated unprotected cytosines to uracil/thymine in both channels, enabling bisulfite-free discrimination of 5hmC versus total methylation (5mC+5hmC) (Fig. 1A). This approach allowed simultaneous quantification of 5hmC (99.5% detection accuracy) and 5mC (via computational subtraction: 5mC+5hmC signal minus 5hmC signal, 98.3% accuracy) at single-cell resolution (fig. S1B), resolving a longstanding limitation of conventional bisulfite-based methods.

**Fig. 1.**
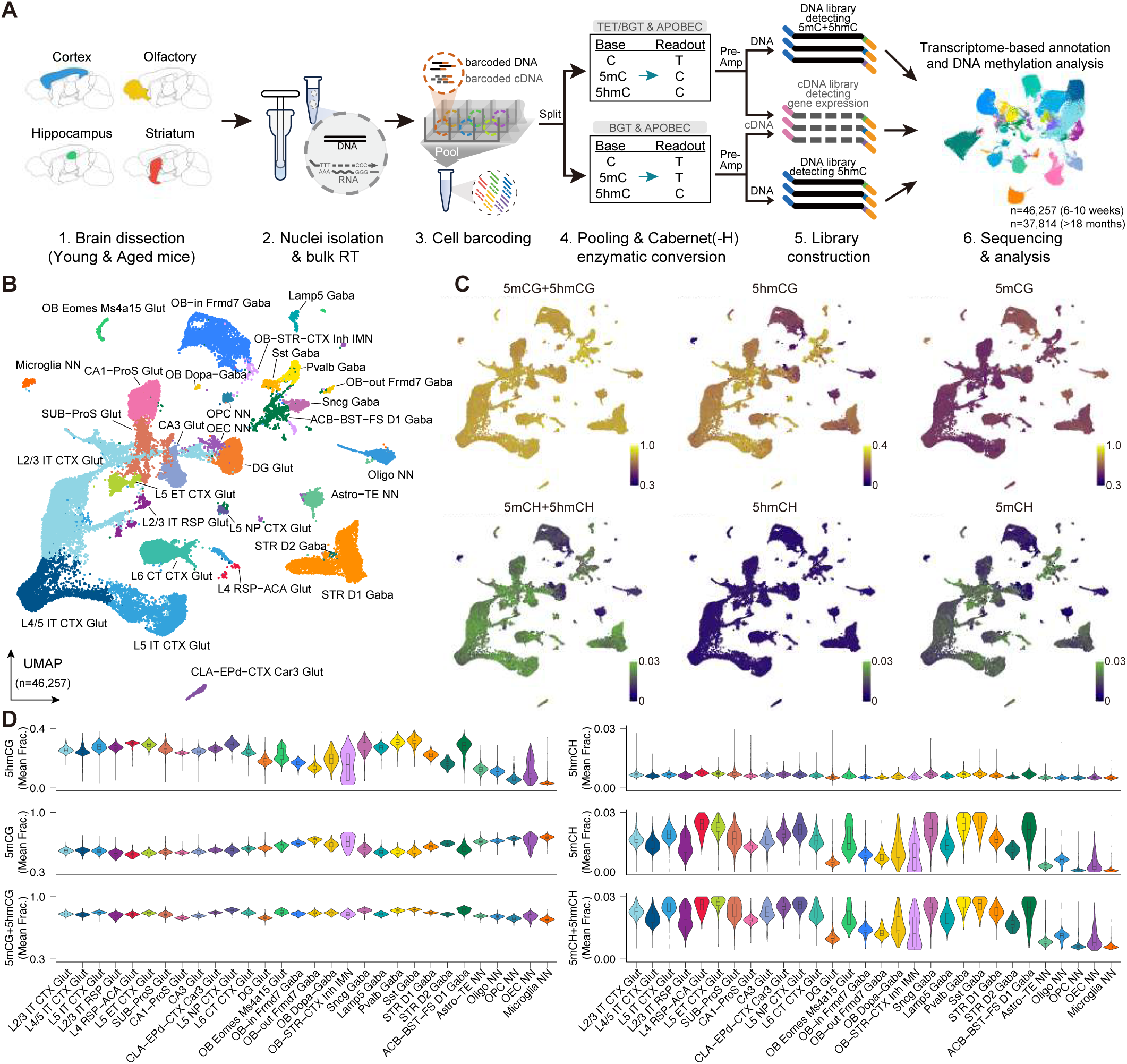
**Multi-modal profiling of mouse brain cells with Joint-Cabernet.** (A) Workflow of brain dissection, nuclei preparation, and joint profiling of RNA, 5hmC, and 5mC. (B) Uniform manifold approximation and projection (UMAP) plot of transcriptome-based clustering of joint profiling data (n=44,608), colored by subclasses annotated with marker genes. (C) Cellular global 5mC+5hmC, 5hmC, and 5mC levels at CG and CH sites, visualized with UMAP. (**D**) Violin plots showing the abundance of 5mC+5hmC, 5hmC, and 5mC levels at CG and CH sites across different cell types.

To synchronize epigenetic and transcriptomic profiles, bulk reverse transcription was performed using a poly-T primer containing a unique reverse adapter different from that used in genomic DNA processing. This design ensured that cDNA and gDNA fragments carried distinct adapter sequences, allowing separate amplification of DNA and RNA libraries. To further distinguish cDNA from gDNA, the reverse transcription reaction incorporated hydroxymethylated dCTP, generating fully methylated cDNA strands. Given the low endogenous mCH (non-CpG) levels in mammalian genomes, this artificial methylation served as a distinct molecular signature (*30*), enabling computational correction of cross-contamination between libraries through mCH ratio quantification.

After cDNA synthesis, both cDNA-RNA hybrid duplexes and genomic DNA underwent synchronized Tn5 transposition with universal transposomes (*31*), followed by identical fragmentation and template-switching workflows to append cell-specific barcodes. This integrative design generated matched 5hmC, 5mC, and transcriptome datasets from identical cells, resolving cell-type-specific methylome-transcriptome relationship analysis while maintaining single-cell resolution.

### DNA methylation and hydroxymethylation landscapes of adult mouse brain

We applied Joint-Cabernet to profile 47,712 nuclei from the adult mouse cortex (superficial and deep layers), hippocampus, striatum, and olfactory regions (6–10 weeks old). Nuclei were labeled with anti-NeuN antibody and sorted via fluorescence-activated cell sorting (FACS), yielding 46,257 RNA-qualified nuclei (Fig. 1A and table S2). For methylation detection, reads with mCH levels <50% were classified as DNA-derived. The following filtering criteria were applied: (i) 5hmC accuracy >98%, (ii) 5mC conversion efficiency (<2% for 5hmC-seq or >96% for 5mC+5hmC-seq), and (iii) unmodified cytosine background <1% (fig. S1, B and C). Ultimately, 44,608 neuronal and non-neuronal cells with matched transcriptomes, 5hmC profiles, and 5mC+5hmC profiles passed quality control for downstream analysis. Sequencing generated an average 8.4 million 5hmC raw reads per cell with 6.0 million aligned reads covering 1.38 million CpG sites. Correspondingly, 8.5 million 5mC+5hmC raw reads were generated per cell, with 5.2 million aligned reads covering 1.45 million CpG sites (fig. S1, D to H). RNA profiling detected on average transcripts from 2,810 genes per neuron and 1,361 genes per non-neuronal cell (fig. S1, I and J).

To enable cell-type-specific methylation analyses, we integrated transcriptomes with a 10x scRNA-seq reference atlas (*2*). Label transfer by comparison to reference datasets identified hierarchical clusters consistent with established neuronal classes and subclasses (fig. S2, A to F). Using marker genes from integrated clusters (table S3), we annotated 13 major types (e.g., excitatory: ‘L2/3 IT CTX Glut’, ‘DG Glut’; inhibitory: ‘STR D1 GABA’) and 30 subtypes (Fig. 1B and fig. S2G). Spatial validation confirmed excitatory neurons resided in their anatomical subregions, while GABAergic subtypes matched tissue origins (e.g., ‘OB-in Frmd7 GABA’ in olfactory, fig. S3A). Cell type annotations predicted by NeuN-based FACS sorting showed high concordance with single-cell transcriptomic clustering across most tissues (fig. S3B). However, in the olfactory bulb, weak NeuN+ fluorescence intensity and overlapping signal distributions between NeuN+ and NeuN− populations resulted in ambiguous gating boundaries. Consequently, FACS-predicted neuronal identities in this region exhibited lower agreement with transcriptional profiles, reflecting biological heterogeneity in NeuN expression levels among olfactory neurons (fig. S3, B to D).

Importantly, global 5hmCG and true 5mCG levels (excluding 5hmCG signals) exhibited pronounced cell-type variability, whereas combined 5mCG+5hmCG signals remained relatively stable across neuronal and non-neuronal populations, demonstrating the importance of resolving 5hmC and 5mC at CG sites (Fig. 1, C and D). In contrast, 5mCH+5hmCH levels (those at non-CpG sites) closely mirrored true 5mCH levels due to sparse 5hmC deposition in CH contexts, suggesting that TET-mediated 5hmC oxidation is highly preferential toward CG motifs.

Consistent with prior studies (*15*, *20*), terminally differentiated neurons exhibited higher global 5hmC levels than non-neuronal cells (Fig. 1D). This accumulation likely reflects the postmitotic state of neurons, in which replication-dependent dilution of 5hmC is minimized. Strikingly, 5hmCG levels varied markedly across neuronal subtypes. GABAergic ‘Sst GABA’ neurons, originating from the medial ganglionic eminence (MGE) and representing the earliest-born cortical interneurons, showed the highest 5hmCG enrichment (mean 31.3% of CG sites). This pattern reflects their early developmental origin and prolonged maintenance in mature circuits. Conversely, dentate gyrus (DG) and olfactory bulb (OB) neurons exhibited reduced 5hmC levels, potentially tied to ongoing neurogenesis in the subgranular (SGZ) and subventricular (SVZ) zones, which replenish these regions with immature, replication-competent cells (*32*, *33*). Notably, single-cell analyses revealed broad intra-subtype variability in 5hmC levels (Fig. 1D), suggesting stochasticity in TET-mediated oxidation—a process where 5mC-to-5hmC conversion rates vary even among transcriptionally similar cells. These findings underscore 5hmC’s role as a cell-type-specific epigenetic regulator and highlight the necessity of direct 5hmC quantification to dissect DNA modification dynamics obscured by conventional bisulfite approaches.

### DNA hydroxymethylation marks actively transcribed DNA

Integrated multi-omic analyses revealed distinct and opposing regulatory roles for 5hmC and 5mC in transcriptional activity. By stratifying genes into non-expressed, low-expressed, and high-expressed categories across neuronal subtypes, we observed pronounced differences in 5mCG and 5hmCG distribution in transcriptionally active loci (Fig. 2A). At transcription start sites (TSS), both modifications were depleted in highly expressed genes, yet the 5hmCG-to-total methylation ratio remained elevated. This suggests that promoter-proximal hydroxymethylation may influence transcriptional initiation. Previous studies reported 5hmC enrichment in active gene bodies of specific neurons and glias (*19*). Here, we demonstrate that all annotated cell types universally exhibit this pattern: 5mCG depletion coupled with 5hmCG accumulation in gene bodies of actively transcribed loci, while combined 5mCG+5hmCG levels remained stable across expression states. This dynamic equilibrium implies that gene body 5hmCG deposition marks transcribed regions, positioning 5hmCG as a more accurate marker for active transcription compared to bisulfite-based CpG measurements. In contrast, CH-context modifications showed minimal 5hmCH (Fig. 2A), explaining why bisulfite-measured gene body CH methylation (primarily 5mCH) correlates more robustly with expression than CpG signals in prior studies (*34*, *35*). Despite the low abundance of 5hmCH, it exhibits a relatively strong positive correlation with gene expression (fig. S4, A and B), but the inherent signal-to-noise limitations restrict the utility of 5hmCH as a robust epigenetic marker. This indicates differential detection of 5hmCG and 5mCG in gene bodies as an accurate epigenetic indicator of transcriptional activity across neuronal and non-neuronal cell types (fig. S4C).

**Fig. 2.**
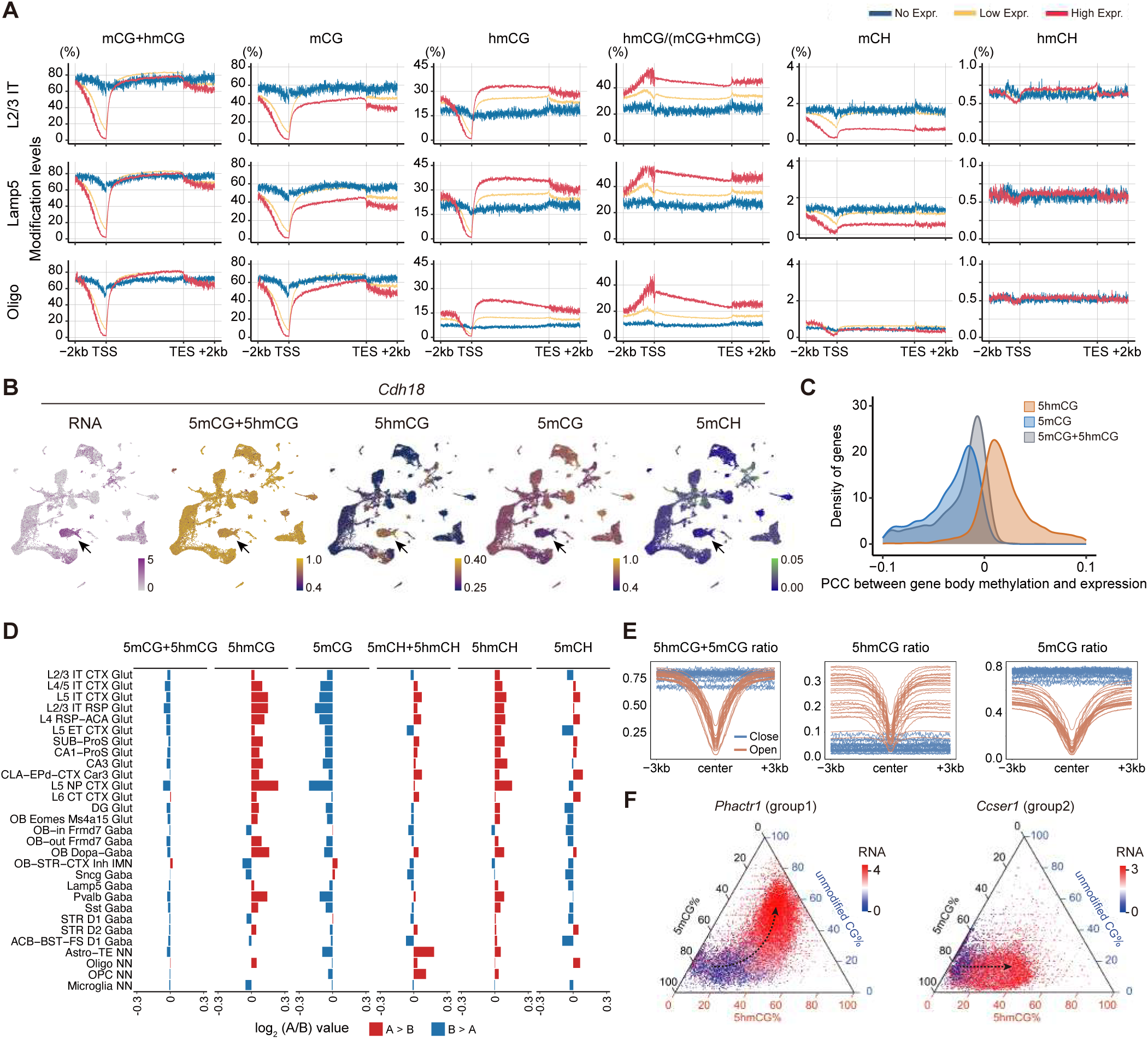
Accurate characterization of the regulatory functions of the DNA methylation and hydroxymethylation. (A) DNA modification levels around genebodies of genes categorized by expression levels: no expression (blue), low expression (yellow), and high expression (red). (**B**) UMAP plots colored by gene expression and genebody modification rates of *Cdh18* (a marker for ‘L6 CT CTX Glut’, which is indicated by a black arrow). (**C**) Distribution of Pearson correlation coefficients (PCCs) between gene expression and genebody CG modifications across single cells. (**D**) Differences in DNA modification levels between A and B compartments at the subclass level. (**E**) DNA modification ratios of 5mCG+5hmCG, 5hmCG, and 5mCG are shown in open or closed chromatin regions, with each subclass represented by one line. (**F**) Ternary plots showing genebody 5mCG, 5hmCG and unmodified CG levels (%), with RNA expression coded in color. Examples from two distinct gene groups are presented to illustrate the differential regulatory mechanisms in gene activation.

For example, *Cdh18*—a marker for layer 6 cortical corticothalamic excitatory (glutamatergic) (‘L6 CT CTX Glut’) neurons—showed reduced 5mCG and elevated 5hmCG in its gene body, correlating with its cell-type-specific transcription, while the 5mCG+5hmCG signal remained relatively stable across different cell types (Fig. 2B). Global Pearson correlation analyses confirmed this trend, with gene body 5hmCG associating positively with expression and 5mCG showing stronger anti-correlation than combined signals (Fig. 2C). Notably, we observed intra-subclass gene activation coincident with 5mCG-to-5hmCG conversion on the gene body, which is undetectable by conventional combined methylation signals (fig. S4D). Together, these findings establish gene body 5hmCG as a dynamic marker of neuronal gene transcription and emphasize the distinct roles of 5hmC and 5mC CpGs in relation to neuronal gene expression.

### Hydroxymethylation is prevalent in active chromatin

In studies of chromatin three-dimensional architecture, it is well established that the genome is partitioned into transcriptionally active A compartments and inactive B compartments (*36*). Using single-cell chromatin conformation data (*6*), we observed that active A compartments are enriched with 5hmC and depleted of 5mC (Fig. 2D and fig. S5A). Analysis of 100-kb-bin compartment score changes (positive scores reflect a shift toward A compartments) across different cell types revealed a positive correlation with 5hmCG levels and a negative correlation with 5mCG levels. However, the combined 5mC+5hmC signal obscures this opposing regulation (fig. S5, B and C), highlighting the importance of distinguishing these modifications. These findings underscore the correlation between cell-type specific gene activity and 5hmC distribution.

Building on previous reports of cell-type-specific CREs (cis-regulatory DNA elements) in the adult mouse brain (*9*), we analyzed the methylation pattern in open chromatin regions. Interestingly, while conventional bisulfite sequencing signals show similar methylation levels at the periphery of open regions compared to closed chromatin regions, the underlying molecular composition differs fundamentally. In contrast, our approach reveals an enrichment of 5hmCG coupled with the depletion of 5mCG at flanking open chromatin centers, a molecular signature invisible to bisulfite-based methods (Fig. 2E and fig. S5, D and E). A previous study demonstrated that MeCP2 binding to 5hmC enhances chromatin accessibility (*19*), which may further contribute to the accumulation of 5hmCG in open chromatin regions through facilitating TET enzyme recruitment. In accordance with that, the heightened accessibility of open chromatin centers may allow TET enzymes to efficiently catalyze oxidative conversion of 5mC to unmodified cytosine, resulting in concurrent depletion of both 5mCG and 5hmCG.

### 5hmCG mediates distinct modes of gene activation in neuronal cells

Certain proteins, such as MeCP2 and MBD3, have been shown to bind directly to 5hmC (*19*, *37*), suggesting that 5hmC may have functional roles beyond being an intermediate in active DNA demethylation. A key question in epigenetic regulation is whether 5hmC itself acts as a distinct regulatory mark. Using Joint-Cabernet data, we can simultaneously examine multiple DNA modifications alongside gene expression, allowing us to systematically assess their interactions and regulatory influences.

To investigate this, we classified genes into distinct groups based on the relationship between gene body modifications and transcriptional activity (fig. S6A). Among 2,644 genes positively associated with 5hmCG, 2,558 (96.7%) exhibited a negative correlation with 5mCG. This strong inverse relationship is consistent with the biochemical conversion of 5mC to 5hmC during active demethylation. However, deeper analysis suggests that demethylation is not the only functional outcome.

Within these actively demethylated and transcriptionally upregulated genes, two major regulatory patterns emerge. Group 1 consists of 1,894 genes that undergo a progressive transition from 5mCG to 5hmCG and subsequently to unmodified CG, accompanied by increasing gene expression across neuronal subtypes (Fig. 2F and fig. S6B). This process occurs in two stages: an initial conversion of 5mCG to 5hmCG, associated with moderate transcriptional activation, followed by a further transition to unmodified CG, linked to a more pronounced activation state.

Group 2, in contrast, consists of 650 genes whose transcriptional activation is primarily associated with the conversion of 5mCG to 5hmCG, without a substantial shift toward unmodified CG (Fig. 2F and fig. S6C). Consequently, bisulfite sequencing, which does not distinguish 5mCG from 5hmCG, fails to detect the dynamics in methylation status for this group. The persistence of 5hmC in this group suggests that 5hmC may have functions independent of its role as a demethylation intermediate, potentially through recognition by specific 5hmC-binding proteins or transcription factors.

Group 1 and Group 2 exhibit divergent genomic and functional characteristics beyond their DNA modification dynamics (fig. S6D). Genes in Group 1 are significantly longer (median length: 221 kb vs. 100 kb in Group 2, p < 2.2e-16) and contain more CpG sites (median CpG number: 1909 vs. 869, p < 2.2e-16). A more complex genomic architecture may favor progressive demethylation, requiring multi-step epigenetic remodeling to achieve transcriptional activation. In contrast, Group 2 genes are shorter with fewer CpGs, suggesting their activation may rely on localized rather than large-scale chromatin restructuring. Notably, while 5hmC is not necessarily a direct driver of gene activation, it has been shown to associate with transcriptionally active chromatin states. Several proteins (e.g., MeCP2) with regulatory potential have been reported to selectively bind 5hmC (*19*), suggesting that 5hmC may serve as a platform for recruiting transcriptional activators or chromatin remodelers.

These gene structural differences align with the functional specialization and expression dynamics of each group. Group 1 is enriched for neurodevelopmental processes (GO: neuron differentiation, synapse assembly, fig. S6A and table S4) and contains 62% of cell-type-specific neuronal markers. Notably, Group 1 genes exhibit significantly higher variability in RNA expression across neuronal subtypes (fig. S6E), reflecting their roles in establishing cell identity through context-dependent regulation. Group 2, however, shows distinct enrichment for basic cellular maintenance functions (GO: organelle organization, primary metabolic process). The lower expression variability of genes in Group 2 across cell types aligns with their housekeeping roles.

Together, the Joint-Cabernet multi-omics approach resolves distinct DNA methylation patterns that would be indistinguishable to conventional methylation analyses. The distribution of 5hmC in Group 2 suggests that 5hmC has functional significance outside of its presence as a demethylation intermediate.

### Differentially hydroxymethylated regions across neurons

Differential 5hmC distribution may play a significant role in neural development and cell differentiation (*38*, *39*). To explore its potential regulatory functions in different neuronal subtypes, we identified 240,150 cell-type-specific differentially hydroxymethylated regions (DHMRs) (Fig. 3, A to C, see Methods). Among these regions, which predominantly overlap with intronic and intergenic elements, 184,573 exhibited hyper-5hmCG in at least one neuronal subtype. Given that over 40% of the DHMRs are located within gene bodies, we investigated their correlation with gene expression patterns across neuronal cell types. By mapping hyper-DHMRs to their corresponding gene loci (table S5), we identified 1,186 genes whose expression levels showed a significant positive correlation with 5hmCG signals within these regions. These DHMRs displayed a marked depletion of 5mCG signals and were also highly cell-type-specific, with minimal variation in the combined 5mCG+5hmCG signals across different cell types (Fig. 3D).

**Fig. 3.**
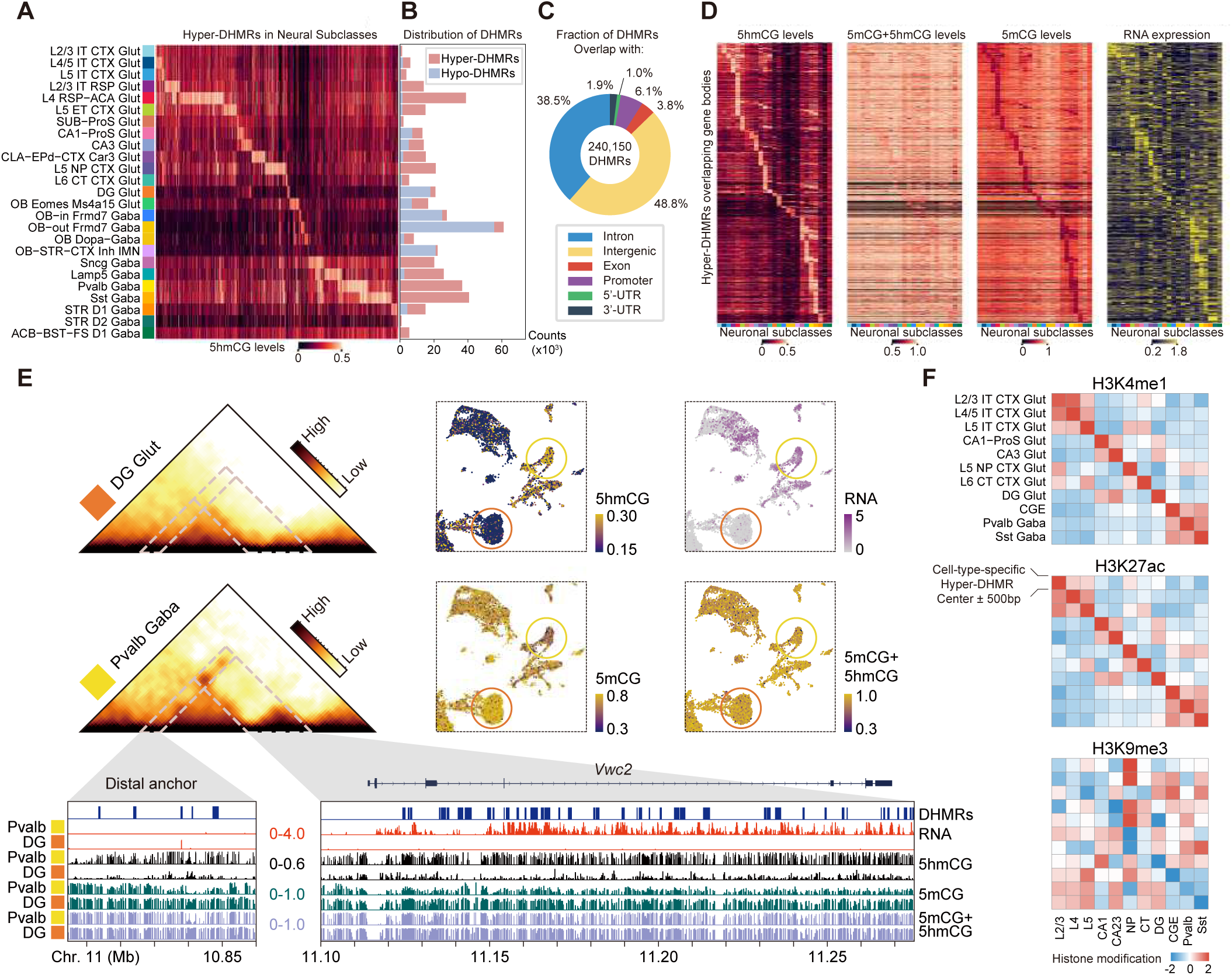
Regulatory roles of differentially hydroxymethylated regions in neurons. (**A**) The 5hmCG ratios in hyper-differentially hydroxymethylated regions (DHMRs) across neuronal subclasses. (**B**) The distribution of DHMRs across neuronal subclasses. (**C**) The annotation of hyper-DHMRs by genomic elements. (**D**) The 5hmCG, 5mCG+5hmCG, and 5mCG levels of genebody hyper-DHMRs, along with the RNA expression levels of the corresponding genes. (**E**) The *Vwc2* distal regulatory region (chr11: 10,805,000-10,860,000), identified by 3C data from Liu et al. (*6*), shows up-regulation of 5hmCG and down-regulation of 5mCG, along with concomitant gene activation in the ‘Pvalb Gaba’ subclass. The contact maps show the interaction intensity. UMAP and IGV visualizations show the *Vwc2* RNA expression levels and the 5hmCG, 5mCG, and 5mCG+5hmCG levels in the upstream distal region. The orange color represents the ‘DG Glut’ subclass, while the yellow color represents the ‘Pvalb Gaba’ subclass. (**F**) Heatmaps showing the average histone modification signals of cell types identified in the Paired-Tag dataset (*42*) on hyper-DHMR centers ± 500 bp. Rows represent the cell-type-specific hyper-DHMRs detected in this study, and columns represent the matched cell types from the Paired-Tag dataset.

In addition to the DHMRs located within gene bodies, we identified a substantial number of DHMRs in intergenic regions, which are likely to play critical roles in gene regulation. To investigate this, we used 3C data to determine the target genes of intergenic regions containing DHMRs (fig. S7A). By focusing on the correlation of interaction strength and gene expression levels (fig. S7B), we identified 3,747 expression-associated interaction anchors (table S6). 5hmCG levels at these distal anchors showed a significant positive correlation with the expression levels of their corresponding target genes (fig. S7C). For instance, the *Vwc2* gene, specifically expressed in ‘Pvalb GABA’ neurons and known for its role in neuronal development in both mouse and human (*40*), showed increased interaction strength with an anchor located 255 kb upstream. This enhanced interaction was associated not only with gene activation but also with a marked accumulation of 5hmCG and significant depletion of 5mCG in the distal anchor (Fig. 3E). Similar results can also be observed for the *Etl4* gene and 160 kb upstream distal anchor in ‘L5 NP CTX Glut’ subclass (fig. S7, D to F).

5hmC has been shown to influence chromatin states by interacting with other epigenetic marks (*41*). Histone modifications, such as H3K27ac and H3K4me1, are known markers of active enhancers, while H3K9me3 is associated with heterochromatin and repressed regions. To investigate the relationship between 5hmC and these modifications, we retrieved histone mark signal data for various cell types from the Paired-Tag dataset (*42*) and analyzed the differential distribution patterns of these marks on cell-type-specific hyper-DHMRs. We found that hyper- DHMRs in each cell type were enriched with active marks, including H3K4me1 and H3K27ac, and showed a depletion of repressive marks, such as H3K9me3 (Fig. 3F).

### Spatially resolved epigenetic landscapes based on location imputation

Precise transcriptional programs underlie the intricate anatomical structures of the mouse brain. While in situ spatial transcriptomics has revealed the diversity of cellular identities and their coordinated functions within brain sections (*43–45*), spatial studies on DNA methylation remain scarce. By integrating high-spatial-resolution MERFISH data (*45*), we inferred the spatial locations of joint RNA and DNA methylation profiles, enabling us to reconstruct spatially resolved DNA methylation landscapes of the mouse brain (Fig. 4A and table S7).

**Fig. 4.**
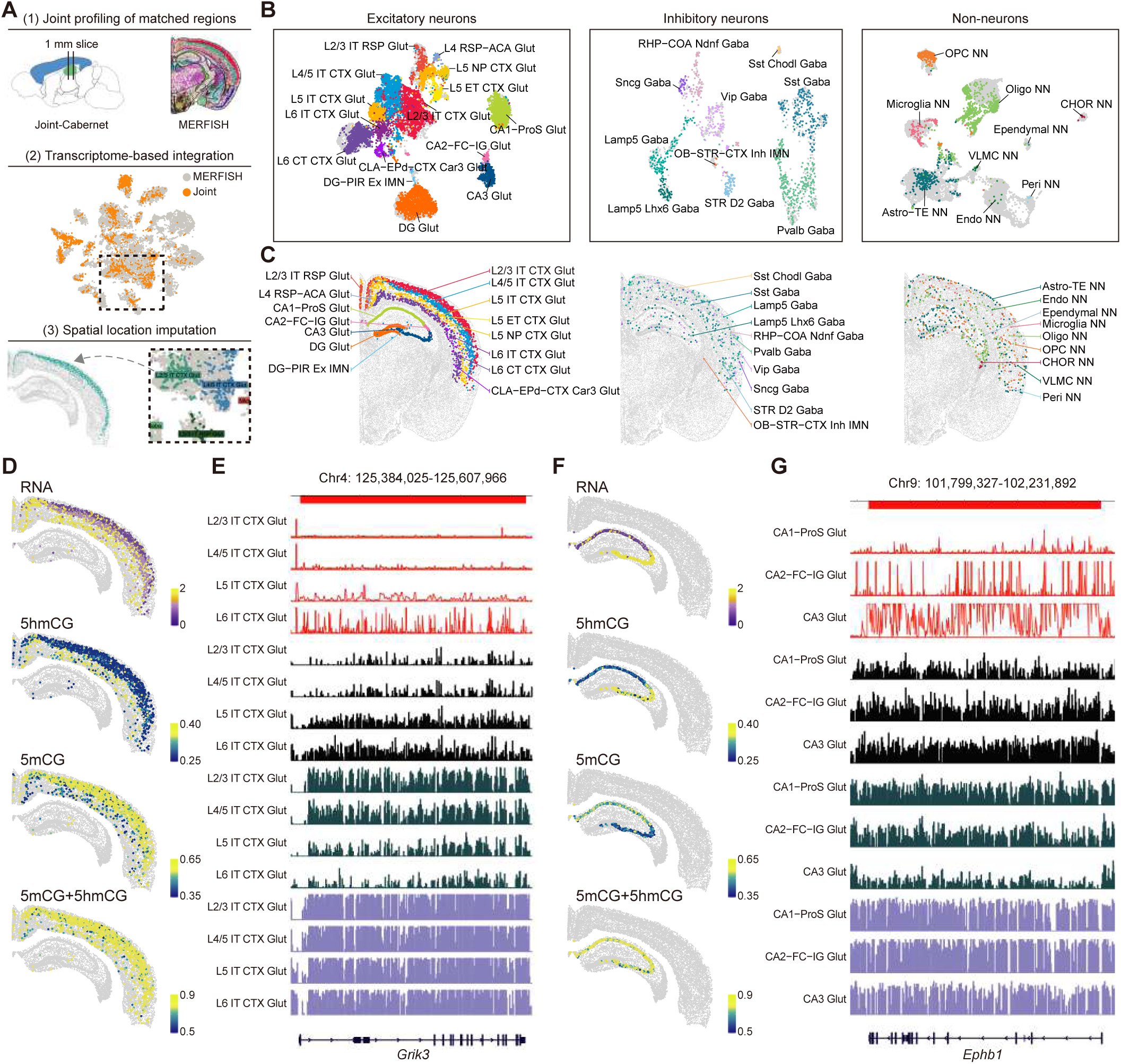
Spatially resolved epigenetic landscapes based on location imputation with MERFISH data. (**A**) Schematic of the spatial projection of 5hmCG and 5mCG profiles through transcriptome-based integration with MERFISH data. (**B**) UMAP plot showing excitatory neurons (left), inhibitory neurons (middle), and non-neurons (right), with gray dots representing cells from the MERFISH data. (**C**) Projected spatial locations of excitatory neurons (left), inhibitory neurons (middle), and non-neurons (right). (**D**) Spatially projected RNA expression, 5hmCG, 5mCG, and 5mCG+5hmCG levels of *Grik3* in related subclasses in the cortex. (**E**) IGV tracks corresponding to **D**. (**F**) Spatially projected RNA expression, 5hmCG, 5mCG, and 5mCG + 5hmCG levels of *Ephb1* in related subclasses in the hippocampus. (**G**) IGV tracks corresponding to **F**.

To ensure the accuracy of spatial location imputation, we performed integration and spatial prediction separately for excitatory neurons, inhibitory neurons, and non-neuronal cells (Fig. 4, B and C). Among these categories, excitatory neurons exhibited pronounced spatial specificity, with many subregion boundaries faithfully preserved in the spatial embeddings. For example, projection neurons corresponding to different cortical layers were accurately localized within their respective layers (‘L2/3 IT CTX Glut’, ‘L5 ET CTX Glut’, ‘L6 CT CTX Glut’).

With spatially resolved 5hmC and 5mC data, we further examined their regulatory functions in a spatial context. For example, the *Grik3* gene, which encodes glutamate receptors involved in various neurophysiological processes (*46*), exhibited a dorsal-ventral gradient of increasing expression in the intratelencephalic (IT) neurons of subcortical layers. This expression pattern was accompanied by a rise in gene body 5hmCG levels and a decrease in 5mCG, with minimal change observed in the combined 5mCG+5hmCG signal (Fig. 4, D and E). Similarly, *Ephb1*, a gene crucial for axon guidance and dendritic spine morphogenesis (*47*), also showed a gradient expression pattern in the cornu ammonis (CA) region of the hippocampus. This gene also demonstrated a positive correlation with 5hmCG and a negative correlation with 5mCG (Fig. 4, F and G). Our analysis revealed that the association of 5hmC with gene activation and 5mC with repression occur in spatially organized patterns, suggesting a contribution to spatial epigenetic regulation.

### Aging-associated DNA modification remodeling of the mouse brain

DNA methylation in mammals changes with aging, a phenomenon often referred to as the ‘methylation clock’. To map age-related DNA modification dynamics, we applied Joint- Cabernet to 37,814 nuclei (table S2) from aged mouse brains (>18 months) and compared them to younger adult counterparts (6–10 weeks). Integrated transcriptomic clustering identified 13 major neuronal and non-neuronal classes (27 subclusters) (Fig. 5A and fig. S8, A and B), with nearly all subclusters exhibiting significant age-associated 5hmCG accumulation (1.06–1.27-fold increase) and concurrent 5mCG depletion (Fig. 5B). Despite this widespread accumulation, the positive correlation between 5hmCG and gene expression was largely preserved in aged cells (fig. S8, C and D). Notably, total methylation levels (5mCG+5hmCG) remained stable at CG sites during aging, masking these opposing dynamics. In contrast, non-CpG context modifications displayed distinct remodeling patterns: cortical neurons showed elevated 5hmCH levels, while olfactory bulb (OB), dentate gyrus (DG), and non-neuronal cells exhibited 5hmCH declines. Conversely, in the 5mC context, 5mCH levels increased universally across cell types (fig. S8E), highlighting context- and cell-type-specific reprogramming during aging.

**Fig. 5.**
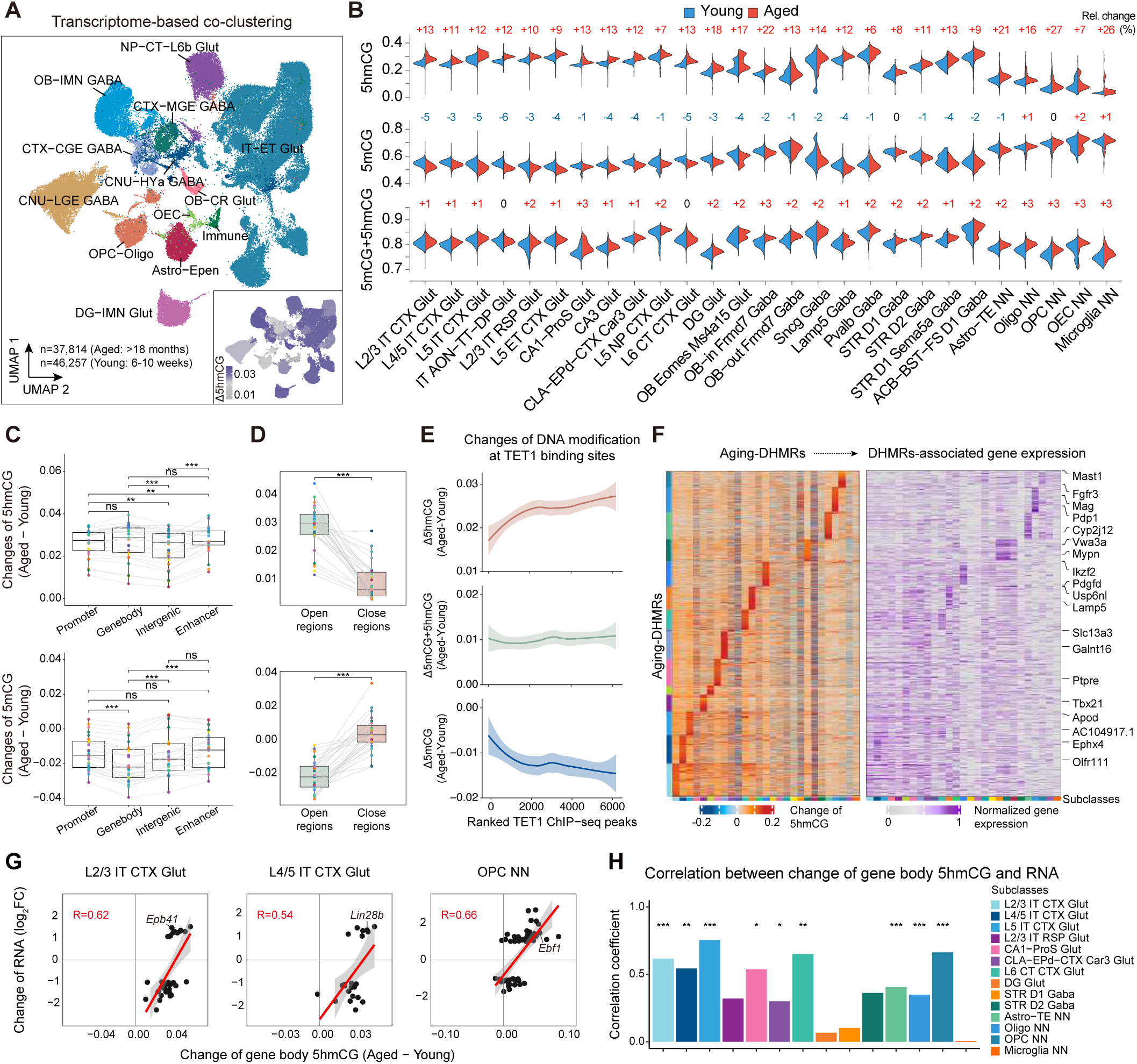
DNA modification changes in aged mouse brain and their associations with RNA expression levels. (**A**) UMAP visualization of transcriptome-based co-clustering for young (6-10 weeks) and aged (18 months) mouse brain cells, colored by class labels. The inset in the right bottom corner shows the increments of 5hmCG levels during aging across subclasses. (**B**) Global DNA modification ratios (5mCG, 5hmCG, and 5mCG+5hmCG) across subclasses demonstrate age-related alterations, with relative rates of change displayed at the top of each comparison. (**C**) The age-related 5hmCG and 5mCG differences in genomic elements. (**D**) The age-related 5hmCG and 5mCG differences in open or closed chromatin regions (right). (**E**) Changes in 5hmCG (brown), 5mCG+5hmCG (green), and 5mCG (blue) levels at TET1 binding sites in the cortex tissue. (**F**) Heatmaps displaying 5hmCG levels (left), and corresponding gene expression (right) in age-associated genebody hyper-DHMRs across subclasses. (**G**) Age-related differences in genebody 5hmCG levels show positive correlations with RNA expression changes for age-related differentially expressed genes (DEGs). (**H**) Pearson correlation coefficients between differences in genebody 5hmCG levels, and log2(Fold-change) of the corresponding RNA expression during aging, across subclasses.

Genomic redistribution of 5hmCG and 5mCG during aging occurred disproportionately in transcriptionally active regions. Gene bodies and enhancers showed preferential 5mCG-to-5hmCG turnover, while closed chromatin domains remained largely unaffected (Fig. 5C). Open chromatin regions exhibited pronounced 5hmCG accrual and 5mCG loss (Fig. 5D), possibly due to facilitated TET-mediated oxidation. This hypothesis is supported by the observation that genomic regions with stronger TET1 binding exhibited more significant 5mCG depletion and 5hmCG gains (Fig. 5E), implicating TET1 as a key driver of age-associated hydroxymethylation. Notably, while closed chromatin regions exhibited less 5hmCG accumulation compared to open regions during aging, we observed significant cell-type-specific differences: excitatory neurons showed markedly higher 5hmCG gains in closed domains than inhibitory neurons and non-neuronal cells (Fig. 5D and fig. S8F). This finding aligns with prior reports of heterochromatin relaxation in aging excitatory neurons (*48*). Strikingly, over 90% of these closed regions with elevated 5hmCG overlapped retrotransposable elements (TEs) (fig. S8G). Given established links between TE reactivation, age-associated inflammation, and neurodegeneration (*49*), our results position 5hmCG as a potential epigenetic biomarker for aberrant TE activation. The preferential hydroxymethylation of TEs in excitatory neurons may reflect their heightened susceptibility to age-related chromatin destabilization and inflammatory cascades.

We identified 392,869 aging-associated hyper-hydroxymethylated regions (aging-DHMRs) across neuronal and glial subclasses, which exhibited striking cell-type specificity in genomic localization (Fig. 5F). Oligodendrocyte-specific aging-DHMRs were enriched at myelination regulators (e.g., *Mag*), while glutamatergic neuronal DHMRs overlapped synaptic signaling hubs (fig. S9A). Functional annotation revealed that aging-DHMR-associated genes were tightly linked to subclass-defining processes: for instance, ‘L6 CT CTX Glut’ neurons showed enrichment for synaptic transmission (GO:0035249), whereas oligodendrocytes were enriched for glial differentiation pathways (GO:0010001). Collectively, these results demonstrate that during aging, genes exhibiting higher cell type-specific expression levels are prone to accumulate 5hmCG, suggesting a transcriptional activity-coupled deposition of this epigenetic mark in a cell type-dependent manner.

To investigate the relationship between 5hmCG accumulation and transcriptional changes, we analyzed aging-related differentially expressed genes (aging-DEGs) from independently validated datasets (*50*). Genes transcriptionally upregulated with age exhibited significantly higher 5hmCG increments across neuronal and glial subtypes compared to downregulated genes (Fig. 5G). Correlation analyses further revealed a positive association between 5hmCG accumulation and age-related gene activation (Fig. 5H). For example, *Ebf1*—a key regulator of oligodendrocyte differentiation and myelination—showed both transcriptional upregulation and pronounced 5hmCG enrichment at its gene body in aged oligodendrocyte precursor cells (OPCs) (Fig. 5G and fig. S9B).

Together, these findings describe the dynamics of age-related methylation changes in the mouse brain. Cell-type-specific 5hmCG accumulation at transcriptionally active loci, possibly driven by TET family recruitment to accessible chromatin, and the coordination between 5hmCG increases and gene upregulation during aging, position 5hmCG as both a biomarker and possible mediator of adaptive epigenetic programming during aging. This occurs without much change in total CG methylation levels, underscoring the necessity of resolving 5hmC and 5mC to decode aging-associated DNA methylation dynamics.

## Discussion

Joint-Cabernet is a single-cell multi-omics platform that resolves long-standing challenges in distinguishing 5hmC and 5mC while synchronously capturing transcriptomes. By applying this method to adult and aged mouse brains, we discovered dynamic hydroxymethylation landscapes that correlate with transcriptional plasticity, chromatin reorganization, and spatial epigenetic patterning. These findings, which would not have been possible by conventional bisulfite sequencing, advance our understanding of epigenetic regulation in brain aging and provide a framework and database for dissecting the role of 5hmC in development and disease.

While Joint-Cabernet achieves high-throughput profiling, its current single-cell genomic coverage remains limited. This constraint necessitates the use of large genomic bins to mitigate sparse data resolution. Additionally, the platform’s transcriptomic profiling exhibits lower gene detection sensitivity compared to dedicated single-cell RNA sequencing methods, requiring integration with reference transcriptomic datasets for robust analysis. Together, these limitations pose challenges in resolving single-cell level correlations and stochastic epigenetic fluctuations, highlighting the need for technical optimizations to enhance sequencing quality.

Joint-Cabernet enables future studies that prioritize defining 5hmC and 5mC dynamics during human brain maturation, aging, and neurodegeneration. We anticipate that human aging—marked by decades of epigenetic remodeling, compared to the shorter lifespan of mice—may exhibit more significant 5hmC accumulation and variability. Such prolonged epigenetic drift could drive transcriptional regulation and dysregulation, potentially correlating with the onset of neurodegenerative diseases. The correlation between 5hmCG accumulation, transcription, and aging may counteract age-related myelin impairment by reinforcing transcriptional programs critical for cellular maintenance. In this way, 5hmCG dynamics could serve as a compensatory mechanism to maintain cellular function during aging, potentially counteracting age-related synaptic and metabolic decline. This hypothesis positions 5hmC as an important epigenetic regulatory mark that may both prevent and contribute to the manifestation of age-related neurodegeneration.

## Methods

### Mouse brain tissues

Experiments involving live mice were approved by the Peking University Laboratory Animal Research Center Institutional Animal Care and Use Committee (IACUC), and all animal experiments were conducted following the ethical guidelines. Adult male C57BL/6J mice aged 7 and 10 weeks were sacrificed, and their brains were rapidly extracted. The superficial cortex, deep cortex, hippocampus, olfactory bulb, and striatum were freshly dissected and immediately placed in ice-cold phosphate-buffered saline (PBS) for subsequent processing. The superficial cortex and deep cortex were further dissected from the cortex by separating them roughly at the midline. For Joint-Cabernet samples, the tissues were dissected and immersed in 1 mL of RNAlater Solution (Invitrogen, AM7020) at 4°C overnight before proceeding to nuclei isolation, fixation, in situ reverse transcription, antibody labelling, and sorting. For samples integrated with the MERFISH data, several P49 mice (three in the first batch and five in the second) were sacrificed to obtain fresh 1000-µm-thick coronal brain sections centered at 7300 µm from the anterior end, as referenced by the Allen Brain Atlas CCFv3 (http://atlas.brain-map.org). These sections were further dissected into left cortex, right cortex, left hippocampus, and right hippocampus before proceeding to nuclei isolation, fixation, in situ reverse transcription, antibody labelling, and sorting.

### Nuclei isolation

Nuclei were isolated as previously described in the Dip-C protocol (*51*), with minor modifications. All buffers used in sample preprocessing were freshly supplemented with RNaseOUT (Thermo Fisher Scientific, 10777019) at a 1:800 ratio. Briefly, brain tissues were first washed in 1 mL of ice-cold PBS and transferred into 2 mL of nuclei isolation medium with Triton (0.25 M sucrose, 25 mM KCl, 5 mM MgCl2, 10 mM HEPES pH 8.0, 1 µM DTT, 0.1% Triton X-100).

Tissues were homogenized using a 2 mL Dounce homogenizer (Sigma D8938) with 5 strokes of the loose (A) pestle and 15 strokes of the tight (B) pestle, followed by centrifugation at 100 g, 4°C for 8 min. The pellet was resuspended in 1 mL of nuclei isolation medium without Triton (0.25 M sucrose, 25 mM KCl, 5 mM MgCl2, 10 mM HEPES pH 8.0, 1 µM DTT) and centrifuged again under the same conditions. The resulting pellet was resuspended in 0.4 mL of nuclei isolation medium without Triton and filtered through a 70 µm strainer (Stemcell, 27216), with two additional washes of the strainer using 0.2 mL of nuclei isolation medium without Triton. Nuclei isolation was completed by another centrifugation at 100 g, 4°C for 8 min, followed by resuspension in 1 mL of PBS-RI.

### Fixation and in situ reverse transcription

Nuclei were fixed and subjected to in situ reverse transcription as previously described in the SPLiT-seq protocol (*52*), with some modifications. Briefly, nuclei pellets were resuspended in 1 mL of ice-cold PBS-RI (PBS supplemented with RNaseOUT at a 1:800 ratio) before adding 3 mL of cold 1.33% formaldehyde solution (16% formaldehyde, Thermo Fisher Scientific, 28906, diluted in 1x PBS) and gently mixed.

The nuclei were fixed for 10 min on ice with occasional inversion for mixing, followed by the addition of 160 µL of 5% Triton X-100 (Sigma-Aldrich, 93443-100 ML) for a 3-min permeabilization on ice. The nuclei were then centrifuged at 500 g, 4°C for 5 min, and resuspended in 500 µL PBS-RI, and mixed with 500 µL of cold 100 mM Tris-HCl pH 8.0 (Invitrogen, AM9855G) and 20 µL 5% Triton X-100. After another centrifugation, the nuclei were resuspended in 200 µL of 0.5x PBS-RI and 4 µL of 5% Triton X-100, then centrifuged and resuspended in 0.5x PBS-RI before counting.

8,000 to 16,000 fixed nuclei per reaction were then in situ reverse transcribed in 20 µL RT reactions (4 µL of 5M Betaine (Sigma-Aldrich, B0300-5VL), 4 µL of 5x RT buffer (Thermo Fisher Scientific), 0.125 µL of RNaseOUT, 0.25 µL of SUPERase-In (Thermo Fisher Scientific, AM2696), 1.43 µL of 7 mM-each dNTP with hydroxymethyl-dCTP instead of dCTP (prepared by diluting dNTPs set (NEB, N0356S, excluding dCTP), and 5-hydroxymethyl-dCTP (Jena Bioscience, NU-932L) in 0.1x TE), 2 µL of 100 µM V2_Poly-dT_TSO+RNA primer (Table S1), 2 µL of Maxima H Minus RTase (Thermo Fisher Scientific, EP0753), 6.195 µL of 0.5x PBS-RI containing the nuclei) with the following program: 50°C 10 min, 3 * [8°C 12 s, 15°C 45 s, 20°C 45 s, 30°C 30 s, 42°C 2 min, 50°C 3 min], 50°C 5 min, and held at 4°C. 30-µL RT mix was then added to the 20-µL reaction for template switching (4.8125 µL of 5M Betaine, 10 µL of 20% Ficoll PM-400 (Sigma-Aldrich, F5415-25 ML), 6 µL of 5x RT buffer, 0.3125 µL of RNaseOUT, 0.625 µL of SUPERase-In, 4 µL of 7 mM-each dNTP with 5-hydroxymethyl-dCTP instead of dCTP, 1.25 µL of 100 µM ME19bp+2G-TSO primer (Table S1), 3 µL of Maxima H Minus RTase), and incubated at 25°C for 30 min and 42°C for 90 min, before being held at 4°C. Every 8 or 16 reactions from the same samples were pooled and 0.45 µL of 10% Triton X-100 was added per reaction, before centrifugation at 1,500 g, 4°C for 15 min. The pellets were resuspended in 1 mL blocking buffer-RI (0.5% BSA in PBS, supplemented with 1:800 volume of RNaseOUT), and again centrifuged at 1,500 g, 4°C for 15 min. All the pellets from each sample were resuspended and pooled in 1 mL blocking buffer-RI before being stored overnight in a 4°C refrigerator.

### Antibody labelling and sorting

Nuclei were antibody labelled as previously described (*26*), with optimizations. To nuclei in 1 mL blocking buffer-RI, 200 µL of antibody mix (1.2 µL (fresh nuclei) or 0.5 µL (fixed nuclei) Alexa Fluor 488-conjugated anti-NeuN antibody (Millipore, MAB377X) diluted in 150 µL PBS-RI and 50 µL blocking buffer-RI and pre-incubated at room temperature for 5 min with protection from light) was added, and incubated for 25 min at 4°C with rotation. DAPI (Sigma, MBD0015) was then added at a final concentration of 2 µg/mL before a 5-min incubation at 4°C with rotation. The nuclei were pelleted through centrifugation at 1,500 g, 4°C for 15 min, and resuspended in 1 mL blocking buffer-RI for a 20-min incubation at 4°C with rotation as a wash step. Finally, the nuclei were centrifuged at 1,500 g, 4°C for 15 min, and resuspended in PBS-RI for single-nucleus sorting into 96-well plates with a BD FACSAria SORP sorter (BD Biosciences) equipped with an 85-µm nozzle. A standard gating strategy based on DAPI staining, SSC/FSC levels, and NeuN antibody staining, was established for all samples of each brain region. Single nuclei were sorted into wells containing 2 µL lysis buffer (20 mM Tris-HCl pH 8.0, 20 mM NaCl, 0.15% Triton X-100, 15 mM DTT, 1 mM EDTA, 3 µg Qiagen Protease (Qiagen, 19157), 0.5 μM carrier ssDNA, and 3 spike-in control (0.1 pg of non-methylated Lambda DNA, 0.03 pg of CpG methylated pUC19, 0.3 pg of fully hydroxymethylated ClaI, as described in Cabernet (*28*), with columns 1–11 sorted with NeuN+ nuclei and column 12 with NeuN- nuclei. After sorting, plates were briefly centrifuged before lysing with the following program: 50°C 1 h, 65°C 1 h, 70°C 15 min and held at 4°C. Lysed plates were stored at -80°C for later use.

### Transposome preparation

The transposon design was inspired by the ‘s3’ design (*29*). The transposon was designed in a unified fashion. These oligos were dissolved in 0.1×TE, mixed at a ratio of 1:1.1, and annealed through heating to 95°C for 1 min and gradual cooling down at a speed of -0.1°C/3 s till 25°C, in 1x annealing buffer (10 mM Tris-HCl pH 8.0, 50 mM NaCl, 1 mM EDTA). The transposase was purified after expression from the pTXB1-Tn5 plasmid (Addgene). Comparable results can be generated with commercial Tn5 transposase (Vazyme Biotech Co., Ltd.). The transposase (6.25 µM monomer) and the annealed transposons (6.25 µM) were then assembled by mixing at a volume ratio of 1:1.1 and incubated at 25°C for 30 min. The assembled transposome was then diluted with Tn5 storage buffer (20% glycerol, 1 mM DTT, 0.1 mM EDTA, 50 mM NaCl, 10 mM Tris-HCl pH 8.0) to the concentration of 300 nM monomer and stored at -80°C. Another kind of Tn5, used for RNA library preparation in the Joint-Cabernet experiments, Nextera i7 Tn5, was also prepared according to the protocol above, except the uracil-containing primer was substituted by the Nextera P7 (Read2) adapter (Table S1).

### Joint-Cabernet library preparation and sequencing

Single-nucleus lysate was transposed in a 10-µL reaction of 10 mM Tris-HCl pH 8.0, 5 mM MgCl2, 8% PEG-8000 (Sigma-Aldrich, P1458), 1.045 nM homemade TSO Tn5 transposome by incubation at 55°C for 10 min. Transposases were removed by addition of 1 μL 2 mg/mL Qiagen protease, combined with 1 μL 12x Tn5 stop buffer of 0.6 M NaCl, 90 mM EDTA, and 0.02% Triton X-100, and incubation at 50°C for 40 min and 70°C for 15 min.

Then, 5.5 µL of contents was moved from each well to a corresponding well in a new plate for the protection and detection of 5mC+5hmC, and the remaining contents in each well was for the protection and detection of 5hmC only.

Gaps before the /U/U/U/ blockage in the transposed DNA were filled, and cell barcodes were added, by the addition of 4.42 μL TSO-gap-PCR mix (2 μL Q5 reaction buffer (NEB), 2 μL Q5 High GC Enhancer (NEB), 0.035 μL 1 M MgCl2, 0.286 µL 5hm-dNTP (7 mM each dNTP, with 5-hydroxymethyl-dCTP instead of dCTP for the plates intended for 5hmC detection) or 5m-dNTP (7 mM each dNTP, with 5-methyl-dCTP instead of dCTP for the plates intended for 5mC+5hmC detection), 0.1 μL Q5 high-fidelity DNA polymerase (NEB, M0491L)), and 0.2 μL of 50 μM well- unique LNA-modified cell barcode TSO primer. The following program was used for PCR amplification: 4°C (pre-cooling of the block), 50°C 3 min, 98°C 30 s, 10 ×[98°C 10 s, 61°C 20 s, 72°C 1 min], 72°C 5 min, held at 4°C.

After cell barcoding, contents from each 96-well plate were pooled together into a single 5-mL tube for purification with the addition of 40 ng sonicated carrier lambda DNA in water and 1270 μL 30% diluted SPRI beads (Beckman Coulter, the dilution buffer was composed of 19% PEG- 8000, 2.5 M NaCl, 10 mM Tris-HCl pH 8.0, 1 mM EDTA, 0.05% Tween 20 (Millipore,655204- 100ML)). The tubes were thoroughly vortexed, incubated at room temperature for 10 min, briefly centrifuged and placed onto magnet racks (Stemcell, 18103) until clear. Four rounds of washing with ∼6.5 mL of 80% ethanol was performed before elution with 48 μL 1 mM Tris-HCl pH 8.0 on a magnet rack for PCR tubes.

The DNA eluted for 5hmC detection was added to a 40-μL BGT reaction mix (1.76 µL 2 mM UDP-glucose (NEB), 8.8 μL NEBuffer 4 (NEB), 4.4 μL T4 Phage β-glucosyltransferase (NEB, M0357L), and 25.04 μL water) before incubation at 37°C for 2 h. The BGT reaction was terminated by addition of 4.4 μL Proteinase K (NEB, P8107S) and incubation at 37°C for 30 min. Purification was performed with 1.8× SPRI beads and the DNA was eluted using 23 μL 1 mM Tris-HCl pH 8.0. DNA denaturation was performed through the addition of 2 μL 0.25 M freshly diluted NaOH, incubation at 50°C for 10 min, and immediate cooling by quickly moving the samples onto an aluminum chill block at 0°C. 25 μL of APOBEC reaction mix (1 μL APOBEC (NEB, E7125L), 5 μL APOBEC reaction buffer (NEB), 0.5 μL BSA (NEB), 1 μL 0.12 M freshly diluted HCl, and 16 μL water) was added while keeping the denatured product on the cold aluminum chill block. Deamination by APOBEC reaction was then performed through incubation at 37°C for 16 h.

The DNA eluted for 5mC+5hmC detection, underwent TET/BGT reaction instead of BGT reaction, which was performed as follows: 33-μL TET reaction mix (2.4 μL water, 18 μL reconstituted TET2 Reaction buffer (NEB), 1.8 μL Oxidation Supplement (NEB), 1.8 μL 100 mM DTT (NEB), 1.8 μL Oxidation Enhancer (NEB), 7.2 μL TET2 (NEB)) was added and mixed before the addition of 9 μL of 1:1250 diluted Fe(II) Solution (NEB) and incubation at 37°C for 1 h, after which the reaction was stopped by the addition of 1.8 μL Stop Reagent (NEB) and incubation at 37°C for 30 min. The TET/BGT reaction products were purified and APOBEC deaminated as described above for the detection of 5hmC, except that the APOBEC reaction time was reduced to 3 h.

Pre-amplification was then performed for both kinds of enzymatic conversion products as follows. After the APOBEC reaction, the first round of pre-amplification was performed by adding 50 μL 2x KAPA HiFi HotStart Uracil+ ReadyMix (Roche, KK2802), 1 μL 100 μM TSO_Read1_primer_V2 and 1 μL 100 μM V2-cDNA-amp-TSO+RNA primer, and running the following PCR program: 98°C 45 s, 14 ×[98°C 15 s, 60°C 30 s, 72°C 3 min), 72°C 3 min, and held at 4°C. The second round of pre-amplification was performed by further adding 1 μL 100 μM TSO_Read2_primer_3U_C/T, and running the following PCR program: 98°C 45 s, 4 ×[98°C 15 s, 60°C 30 s, 72°C 3 min), 72°C 3 min, and held at 4°C. The pre-amplification products were purified using 1.6x SPRI beads and eluted in 52 μL of EB, with 24 μL of which headed for library preparation of 5-(hydroxy)methyl-cytosines detection, and another 24 μL of which headed for library preparation of RNA detection.

Libraries detecting 5-(hydroxy)methyl-cytosines were both prepared by adding 25 μL 2x KAPA HiFi HotStart ReadyMix (Roche, KK2602), 0.5 μL 100 μM TSO custom P5 index primer and 0.5 μL 100 μM TSO custom P7 index primer to the 24 μL pre-amplified product, and running the following PCR program: 98°C 45 s, 5 ×[98°C 15 s, 60°C 30 s, 72°C 90 s), 72°C 3 min, and held at 4°C. Purification and size-selection were performed respectively with DNA Clean & Concentrator-5 (ZYMO, D4013) and Select-A-size DNA Clean & Concentrator (ZYMO, D4080, cutoff chosen at 150 bp) according to the manufacturer’s guidelines.

For RNA library preparation, 24 µL of pre-amplified product from each enzymatic conversion path was pooled with its counterpart originating from the same source plate. For example, plate A was split into plate A1 (for 5mC+5hmC detection) and plate A2 (for 5hmC detection), which were cell barcoded, pooled, purified, enzymatically converted in different ways, and pre-amplified as 2 conjugated samples, according to the methods described above; the pre-amplified products from plate A1 and A2, which both represent gene expression profiles of cells in plate A, were pooled to generate a combined 48-μL (24-µL each) product for subsequent RNA library preparation.

The combined 48-μL pre-amplified products were added 50 μL 2x KAPA HiFi HotStart ReadyMix, 1 µL 100 μM TSO_Read1_primer_V2 and 1 μL 100 μM V2-cDNA-amp-TSO+RNA primer before running the following PCR program: 98°C 45 s, 5 ×[98°C 15 s, 60°C 30 s, 72°C 3 min), 72°C 3 min, and held at 4°C. This PCR product was purified using 1.2x SPRI beads, and eluted in 100 μL EB. The elution was termed as cDNA Amp product, as it had been further enriched for RNA reads.

4 ng of the cDNA Amp product was transposed in a 10-µL reaction of 10 mM Tris-HCl pH 8.0, 5 mM MgCl2, 8% PEG-8000, 10.8 nM home-made Nextera i7 Tn5 transposome by incubation at 55°C for 10 min, and stopped in a 12-μL reaction as described above. 37 μL of Q5 mix (15.43 μL water, 10 μL Q5 reaction buffer (NEB), 10 μL Q5 High GC Enhancer (NEB), 0.07 μL 1 M MgCl2, 1 μL dNTP mix (NEB, N0447S, 10 mM each), 0.5 µL Q5 high-fidelity DNA polymerase), 0.5 μL 100 μM TSO custom P5 index primer and 0.5 µL 100 μM Nextera P7 index primer, were added before performing the following PCR procedures: 65°C 3 min, 98°C 30 s, 8 ×[98°C 10 s, 60°C 30 s, 72°C 90 s], 72°C 2 min, and held at 4°C. The PCR products were added 54 μL EB and 56 μL SPRI beads for removal of the over-long fragments by retaining 150 μL of the supernatants, which were further added 20 μL SPRI beads, purified, and eluted by adding 20.5 μL EB. 20 μL of the elution was then collected as crude RNA library, which was pooled and further size-selected using Select-A-size DNA Clean & Concentrator (cutoff chosen at 300 bp) to get the final pooled RNA libraries.

We designed a custom balance library for improving the low nucleotide diversity caused by the fixed Tn5 ME 19 bp sequence following the cell barcode in the Read1 and the low-CG nature of (hydroxy)methyl-libraries that convert unmodified Cs to Ts (Table S1). It must be noted that custom sequencing primers were used for Illumina sequencing as required by the library structures. For sequencing that only involved the (hydroxy)methyl-libraries, all sequencing primers were replaced by the custom sequencing primers, while in the cases involving sequencing the RNA libraries, custom sequencing primers related to the P7 end were spiked in, as the RNA libraries had standard Nextera P7 ends. Libraries were first sequenced in a 2 ×150 bp mode on an Illumina MiSeq system for QC, with the TSO custom balance library added at a ratio of 30%. The libraries passing QC were sequenced on an Illumina NovaSeq 6000 system in a 2 × 150 bp mode, using one S4 flow cell for around 24 (hydroxy)methyl-libraries or 288 RNA libraries, with the TSO custom balance library added at a ratio of 20%.

### Quality control

Raw sequencing reads of DNA and RNA for each single cell were processed following the Joint- Cabernet pipeline (Code is available on GitHub at: https://github.com/yunlongcaolab/MouseBrain_epiAtlas.git). Individual cell libraries were filtered as follows: DNA 5mC+5hmC libraries require align rate >20%, coverage >0.001, pUC19 mCG ratio >0.96, lambda mCG ratio <0.01; DNA 5hmC libraries require align rate >20%, coverage >0.001, pUC19 mCG ratio <0.02, lambda mCG ratio <0.01, ClaI mCG ratio >0.98; RNA libraries require align rate ≥20%, separated RNA reads ratio ≥70%, RNA reads ≥20,000, gene number ≥300, Qualimap Intronic ≥40%, Qualimap Intergenic ≤30%.

### Transcriptome-based integration

We applied the Robust Principal Component Analysis (RPCA) method in Seurat (v5.1.0, https://github.com/satijalab/seurat) (*53*) to integrate snRNA-seq data from multiple brain regions, including the cortex, hippocampus, olfactory bulbs, and striatum, with the reference dataset from Yao et al. (*2*). The ’RunPCA’ function was utilized to compute the top 20 principal components (PCs) based on variably expressed genes. Clustering was conducted on the integrated expression data using a shared-nearest-neighbor (SNN) graph-based approach with Louvain community detection, implemented through the ’FindClusters’ function with a resolution of 0.8, before visualization with Uniform Manifold Approximation and Projection (UMAP) (*54*). After integration, each cell in a Seurat cluster was assigned the most prevalent original label of the cells from the reference dataset in that cluster.

### DNA methylation around gene bodies with different expression levels

The median value of non-zero expression levels in the single-cell expression matrix from the reference dataset from Yao et al. (*2*) was calculated, and genes were then divided into 3 groups according to their average expression levels in the reference dataset for each subclass: genes with zero expression, genes with expression no larger than the median, and genes with expression over the median. The average methylation levels around the gene bodies of the genes in the 3 groups were then calculated by applying the ‘computeMatrix’ function from deeptools (*55*) onto the subclass-level methylation BigWig files generated from ALLC files, with the following parameters: regionBodyLength = 5000, binsize = 5, upstream = 2000, and downstream = 2000, before plotting using the ‘geom_line’ function from the ggplot2 package (https://ggplot2.tidyverse.org) in R.

### Correlation between RNA expression and gene body DNA methylation

We calculated the Pearson correlation coefficients (PCCs) between RNA expression levels and gene body mean DNA methylation levels across cells, for all genes, with the p-values adjusted using the BH method. To estimate false positives due to randomness, we also performed 5000 random shuffles of the samples and calculated the PCCs for a negative control.

The ‘geom_density_line’ function from the R package ggplot2 was used to plot the density distribution of PCCs. The ‘geom_pointdensity’ function from ggplot2 was utilized to create scatter plots depicting the relationships between RNA expression and DNA methylation levels, with fitting lines generated by the ‘stat_cor’ function.

### 3D-compartment-level analysis of DNA modifications

We performed a comprehensive analysis of Hi-C data for different subclasses by applying the cooler package (v0.10.3, https://github.com/open2c/cooler) (*56*) onto the 100-kb-resolution contact matrix for each subclass from Liu et al. (*6*). The Hi-C matrices were normalized using the cooler balance tool. We then converted the processed data from COOL format to HDF5 format using hicConvertFormat (v3.0) (*57*), enabling efficient downstream analysis. To examine the structural variation of the contact matrix, we performed principal component analysis (PCA) using the hicPCA (*57*) tool. The first three principal components (PC1, PC2, and PC3) were extracted and saved in BEDGraph format. For further analysis, we classified the genome into two compartments (A and B) based on the sign of the PC1 values, and we filtered out any entries where PC1 was zero. Finally, we calculated the 5hmC, 5mC, and 5mC+5hmC levels in CG and CH contexts across A/B compartments for each subclass.

### Chromatin accessibility and DNA methylation dynamics

We obtained publicly available subclass-specific ATAC-seq chromatin regulatory elements (CREs) (*9*). Open chromatin regions were defined as genomic intervals centered at CRE midpoints and extending 3 kb upstream and downstream. Closed chromatin regions were designated as consecutive 6 kb genomic intervals devoid of any CRE overlap. We then quantified the levels of 5hmCG, 5mCG, and the combined 5mCG+5hmCG ratio in both open and closed chromatin regions across different subclasses.

### D(H)MR analysis

First, we merged single-cell ALLC files into pseudo-bulk ALLC files according to the labels from clustering analysis, using the ‘merge-allc’ command from ALLCools (v1.0.8, https://github.com/lhqing/ALLCools). Next, we divided the genome into homogeneous methylated candidate regions for DMR-calling (‘segments’), using the ‘segment --min_cpg 3 -- max_bp 2000’ command from wgbstools (v0.2.0, https://github.com/nloyfer/wgbs_tools) with the subclass-level merged ALLC files. The mean CG methylation levels for the segments in individual cells were then generated with the ‘generate-dataset’ command from ALLCools. We then conducted DMR-calling in 3 stratified levels: (1) among excitatory neurons, inhibitory neurons, and non-neurons; (2) among all neuronal subclasses; (3) among all non-neuron subclasses. Segments with lengths of at least 200 bp then underwent DMR-calling using two-sided Wilcoxon rank-sum test by comparing cells from 1 group and cells from all other groups. Segments were regarded as DMRs if the BH-adjusted p-values were not above 0.05, the numbers of cells with non-NA values in the group were at least 10, and the mean methylation levels of the group differed from the robust means of all groups (the mean methylation levels of groups with methylation levels between 25th and 75th percentiles) by over +0.3 (hyper) or below -0.3 (hypo). The same process was carried out for DHMR analysis, except that the threshold of the differences of mean methylation levels from robust means were changed from +/-0.3 to +/-0.2 as the dynamic range of 5hmCG levels is smaller than 5mCG+5hmCG levels. Annotation of the DMRs and DHMRs was carried out using the ‘annotate_by_beds’ method of the ‘RegionDS’ class from ALLCools with the parameter ‘slop = 250’. For overlapping of DMRs and DHMRs with gene bodies and promoters, only genes with expression levels over 0.2 in at least 1 subclass and the coefficient of variation (CV) of expression levels among subclasses over 0.5, were used, and the overlapping thresholds were set to max(0.6*min(gene_length, D(H)MR_length), 500 bp (for overlapping gene bodies) / 100 bp (for overlapping promoters)).

### Linking distal regions with hyper-DHMR to genes through 3C data

First, loops (dots) were called using matching neuronal subclass pseudo-bulk imputed 10-kb-resolution 3C interaction matrixes generated through snm3C-seq (*6*) with the command ‘chromosight detect --min-dist 20000 --max-dist 500000’ from chromosight (*58*). Next, only loops with one anchor overlapping genes with expression levels over 0.2 in at least 1 subclass and the coefficient of variation (CV) of expression levels among subclasses over 0.5 (the genic anchor), and the other anchor not overlapping with any genes but having at least 1 hyper-DHMR within the 30-kb regions centering at this anchor (the intergenic anchor with hyper-DHMR), were preserved, linking distal regions with hyper-DHMR to genes. The 5hmCG levels of the 30-kb regions centering at the intergenic anchor with hyper-DHMR were then calculated through the ‘generate-dataset’ command from ALLCools. The 3C interaction between the intergenic anchors and the linked genes were calculated by summing up the balanced 3C interaction between the intergenic anchor and all the 10-kb bins overlapping with the gene. The Pearson correlation coefficients (PCCs) between the RNA expression levels of the linked genes and the 5hmCG levels in the distal regions, as well as the Pearson correlation coefficients (PCCs) between the RNA expression levels of the linked genes and the 3C interaction between the intergenic anchors and the linked genes, were then calculated.

### Histone modification analysis in hyper-DHMRs

We analyzed the enrichment of histone modifications in the hyper-DHMRs across different subclasses to explore their potential regulatory roles. The histone modifications examined included H3K4me1, H3K27ac, and H3K9me3. Using publicly available subclass-specific histone modification data (GSE152020), we quantified the signal intensity of these modifications in hyper- DHMRs.

### Spatial projection of Joint-Cabernet cells using MERFISH data

We selected cells with z-values ranging from 7.33 to 7.34 and located in the brain regions of isocortex and hippocampus in the downloaded MERFISH data (*45*) as reference data for spatial projection, and categorized the cells into excitatory neurons, inhibitory neurons, and non-neurons according to the ‘subclass_transfer’ information in the metadata. Accordingly, we categorized our Joint-Cabernet data of brain coronal sections into excitatory neurons, inhibitory neurons, and non-neurons by integrating with 10x RNA-seq data from Yao et al. (*2*). The excitatory neurons, inhibitory neurons, and non-neuronal cells from those two sources were then integrated separately before performing label transfer. Spatial projection was then carried out by assigning the coordinates of the nearest MERFISH cell in UMAP coordinates with matching brain region and subclass, to Joint-Cabernet cells.

### Aging-related clustering and D(H)MR analysis

Integration of Joint-Cabernet RNA data from both young and aged mice with the downsampled 10x RNA-seq data from Yao et al. (*2*), was performed as mentioned above using Seurat, except that we used the intersection of the features found by ‘SelectIntegrationFeatures’ and the top 1000 subclass-specific marker genes identified during the prior integration of young mouse Joint- Cabernet RNA data with the dataset from Yao et al. The D(H)MR analysis was conducted as mentioned above, except that the comparisons were made between the cells from young mice and the cells from aged mice in the same subclass, instead of between one subclass and the other subclasses, and that the requirement for absolute differences in modification rates was adjusted to >0.05.

### DNA methylation levels on TET1 ChIP-seq peaks

To investigate the relationship between TET1 occupancy and DNA methylation changes during aging, we analyzed publicly available TET1 ChIP-seq data (GSE108768). Peak regions were ranked based on signal intensity, and DNA methylation levels were calculated across these regions. Specifically, we compared the aging-related changes in 5hmCG, 5mCG, and 5mCG+5hmCG levels along the gradient of increasing TET1 ChIP-seq signal.

### Data and materials availability

Raw and processed data of Joint-Cabernet were deposited under the accession number GSE294517. The analysis code is available at https://github.com/yunlongcaolab/MouseBrain_epiAtlas.git.

## Acknowledgments

This work is financially supported by National Natural Science Foundation of China (2023011477, Y.C.) and Changping Laboratory (2025B-06-01, Y.C.). We thank Y. Chen and C. Geng at Peking University High-throughput Sequencing Center operated by BIOPIC for sorting of nuclei and assisting with Illumina Miseq sequencing. We used AI-assisted tools, including ChatGPT and DeepSeek, to help edit and polish the manuscript for language clarity.

## Author contributions

Y.C. and X.S.X. supervised the study. T.Y., Y.B., L.R., Y.H., and Y.C. developed the Joint- Cabernet. Y.B., T.Y., L.R., Y.H., F.Y., Y.L., A.Z., Y.L., T.T., N.K., D.C., X.X., Q.W., W.C., and Y.Z. generated and preprocessed the single-cell data. Y.B., T.Y., L.R., Y.H., F.Y., A.Z., and Y.L. performed bioinformatic analyses. Y.B., T.Y., and Y.C. wrote the manuscript. All authors edited and approved the manuscript.

## Supplementary Figures

**Fig. S1.**
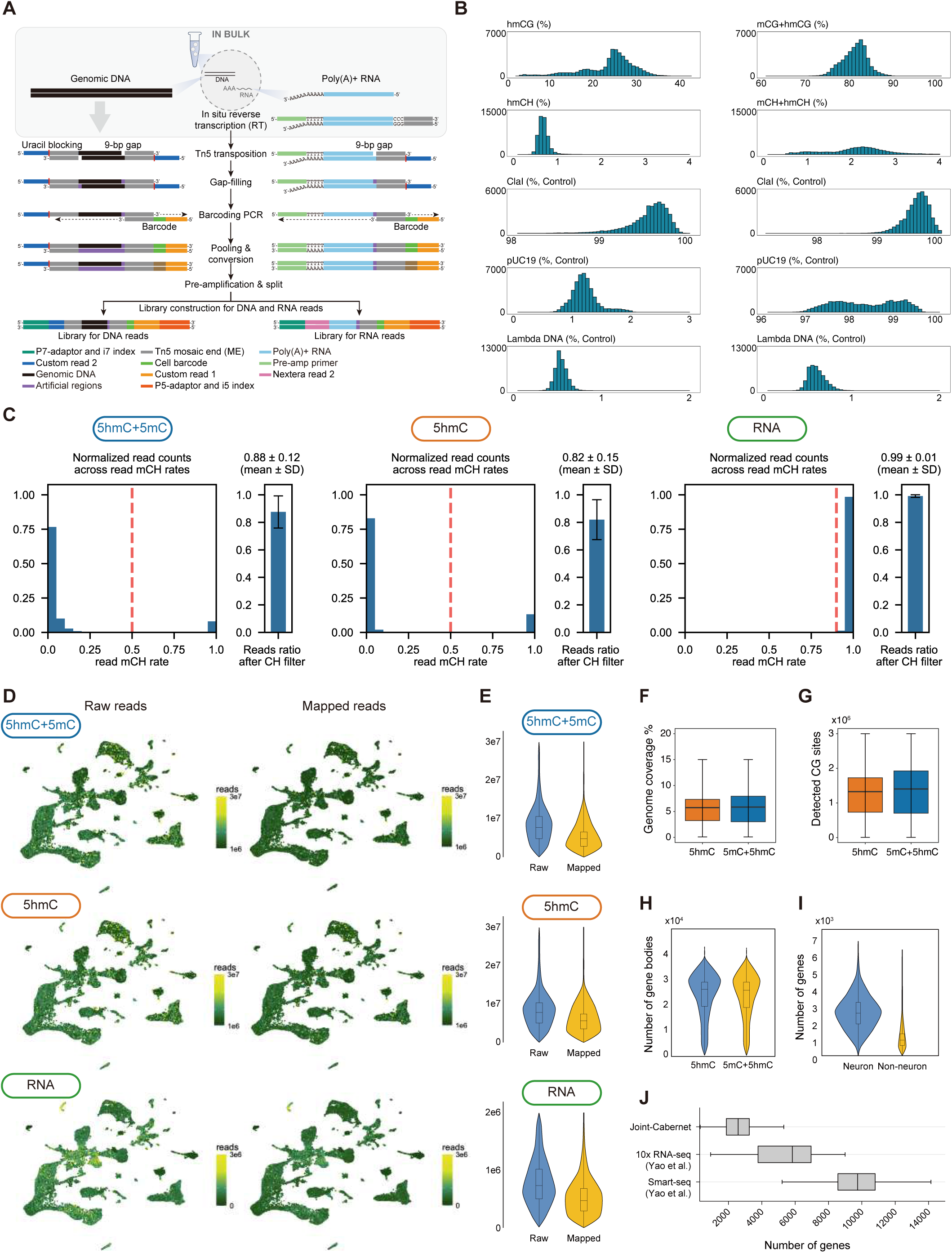
Overview of joint RNA, 5hmC, and 5mC profiling and data quality assessment. (**A**) Schematic of joint profiling of RNA, 5hmC and 5mC. (**B**) Distribution of the 5hmC and 5mC+5hmC levels at genomic CG, CH sites, and spike-in controls (5hmC control, ClaI; 5mC control, pUC19; unmodified C control, lambda DNA). (**C**) The fractions of the filtered reads based on mCH rate of each read. In the 5mC+5hmC and 5hmC libraries, reads with at least 3 CH sites and read-level mCH ratio < 0.5 were assigned to DNA. In the RNA library, reads with at least 3 CH sites and read-level mCH ratio ≥ 0.9 were assigned to RNA. (**D, E**) Distribution of the numbers of raw reads and aligned reads in this study shown by UMAP and violin plots. (**F, G**) Distribution of genomic coverage and number of CpG sites detected in each cell. (**H**) Distribution of the numbers of gene locus detected per cell in the 5hmC profiling and the 5mC+5hmC profiling. (**I**) Distribution of the number of detected genes per neuronal and non-neuronal cell, in the RNA profiling. (**J**) Boxplot showing the numbers of genes detected in each cell by different single-cell RNA-seq approaches.

**Fig. S2.**
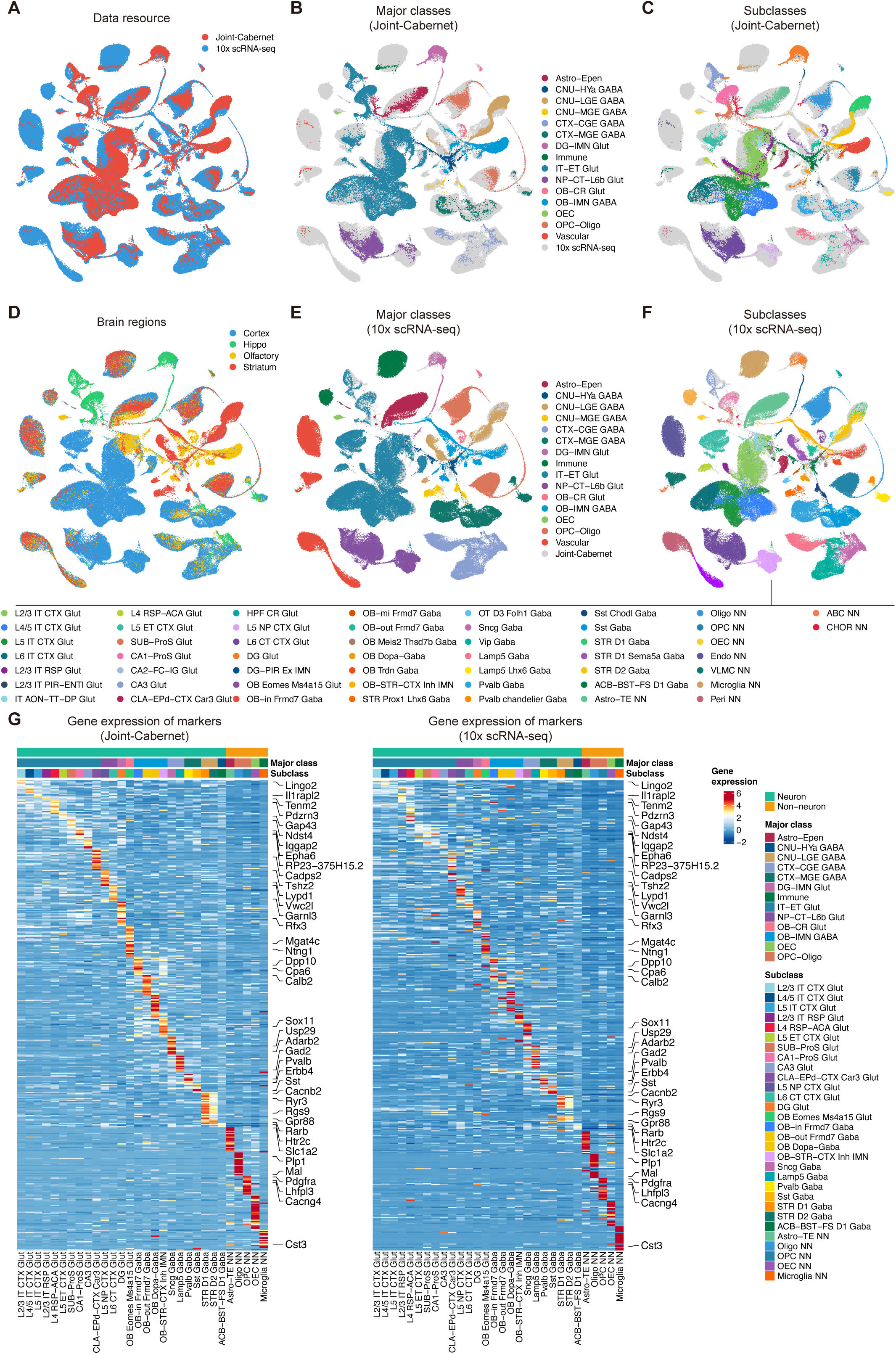
Integration of joint-Cabernet and 10x scRNA-seq data (**A-C**) Transcriptome-based integration UMAP colored by datasets, major classes and subclasses of joint profiling data in this study. (**D-F**) Integration UMAP colored by dissection regions, major classes and subclasses of the 10x scRNA-seq dataset. (**G**) The expression levels of marker genes in this study and the previously published 10x scRNA-seq dataset.

**Fig. S3.**
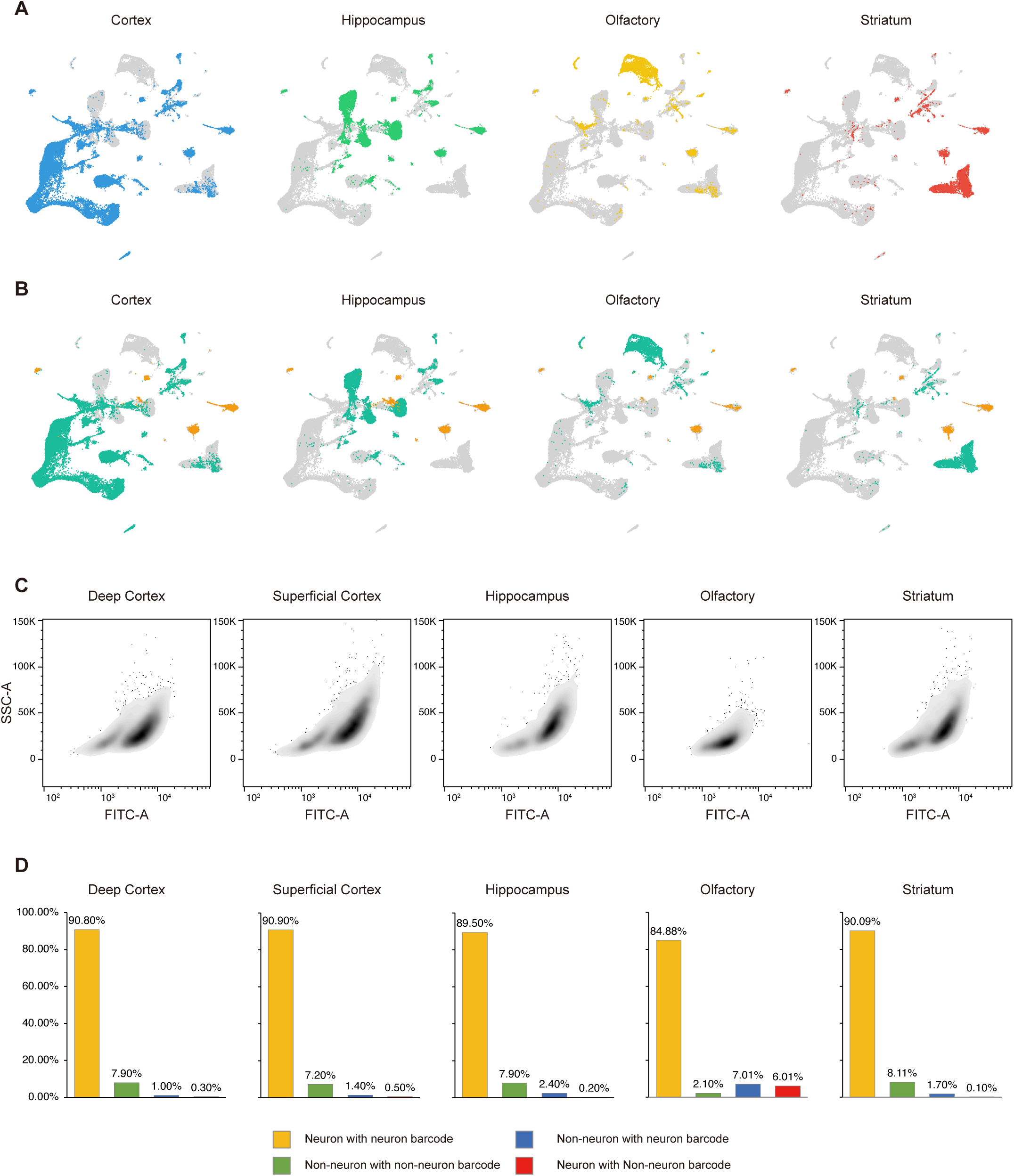
Accuracy of FACS (**A**) UMAP colored by dissection regions. (**B**) UMAP colored by neuron or non-neuron labels. (**C**) Representative distribution of FACS signals with samples from different dissection regions. The FITC-A signals represent the NeuN antibody signals. (**D**) The accuracies of FACS labeling compared with annotated neuron or non-neuron labels.

**Fig. S4.**
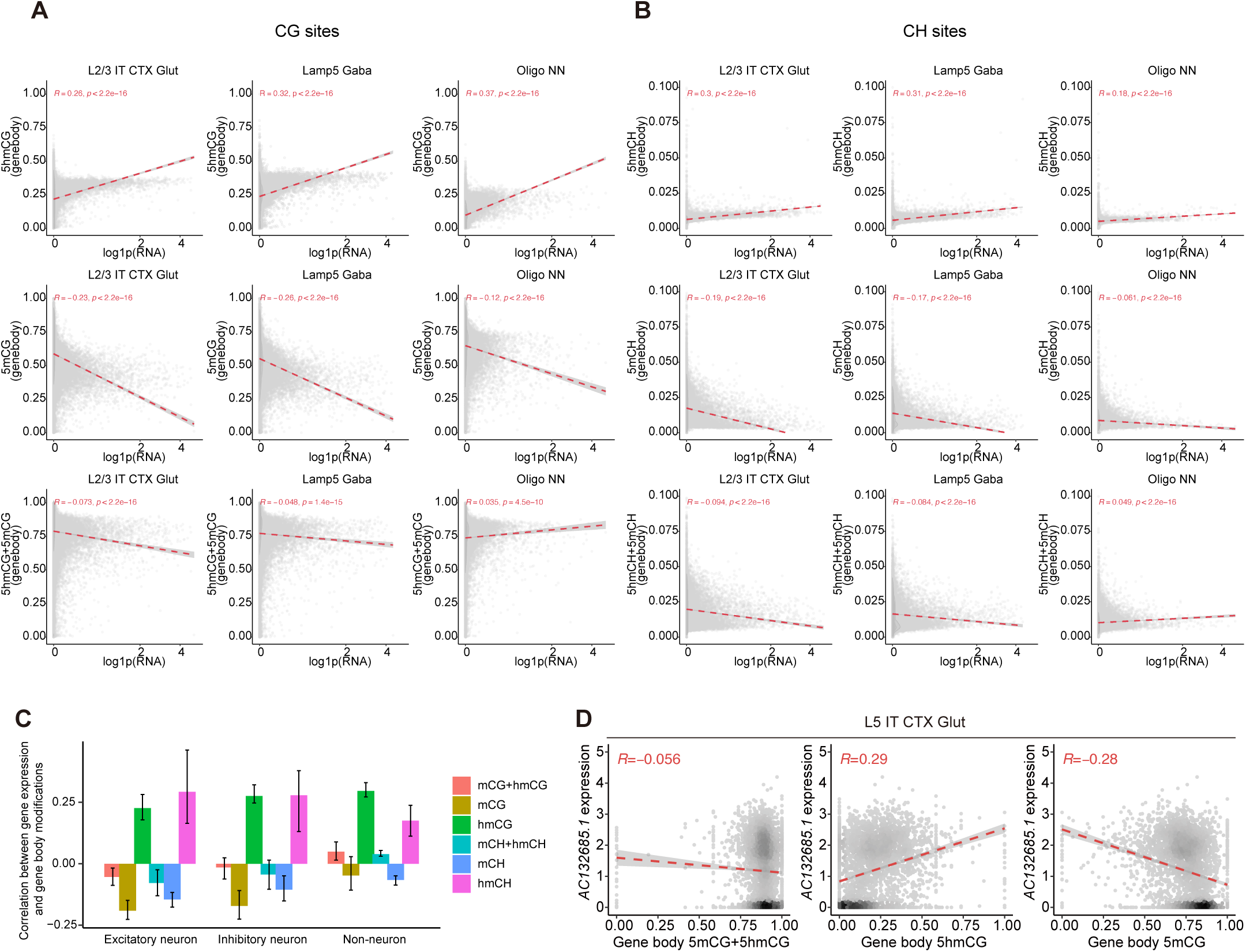
Relationship between DNA modifications and gene expression across genomic contexts (**A, B**) Scatter plots showing the relationship of 5mC+5hmC levels, 5mC levels, 5hmC levels with gene expression levels at CG and CH contexts in genebodies of genes in different subclasses. (**C**) Distribution of Pearson correlation coefficients (PCCs) between gene expression and genebody 5mC+5hmC, 5mC, and 5hmC modifications at CG and CH sites contexts in genebodies of genes in different subclasses, with the subclasses categorized into excitatory neurons, inhibitory neurons and non-neurons. The PCCs corresponds to the R values in **A** and **B**. (**D**) Scatter plots showing the relationships between *AC132685.1* expression and genebody modifications in single cells of the ‘L5 IT CTX Glut’ subclass.

**Fig. S5.**
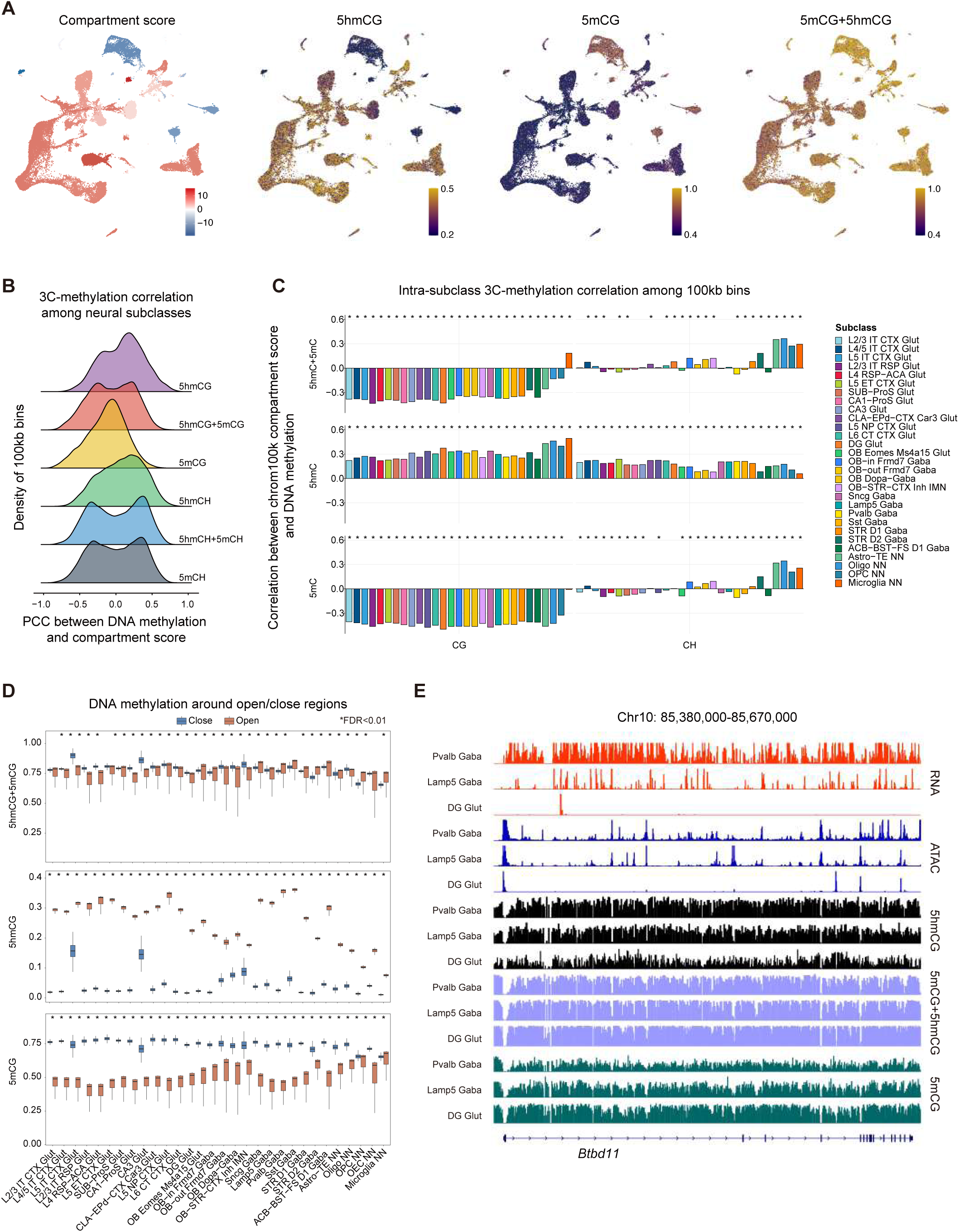
DNA modifications in relation to chromatin accessibility and chromatin conformations across cell types. (**A**) UMAP showing compartment scores and DNA modification ratios across cells from different subclasses, highlighting the association between DNA modification patterns and chromatin compartmentalization on the 100-kb bin of chr8: 17,100,000-17,200,000. (**B**) Distribution of PCC values between compartment scores and 100-kb-bin modification ratios among neural subclasses. (**C**) Intra-subclass PCC values between compartment scores and modification ratios among 100- kb bins. (**D**) Boxplots depicting DNA modification levels around open or closed regions for neuronal and non-neuronal subclasses. (* indicates FDR < 0.01.) (**E**) IGV tracks of a genomic locus showing the distribution of RNA expression, ATAC-seq signals, and DNA modification patterns for various cell types, revealing coordinated changes in gene expression, chromatin accessibility and multiple DNA modifications at specific loci.

**Fig. S6.**
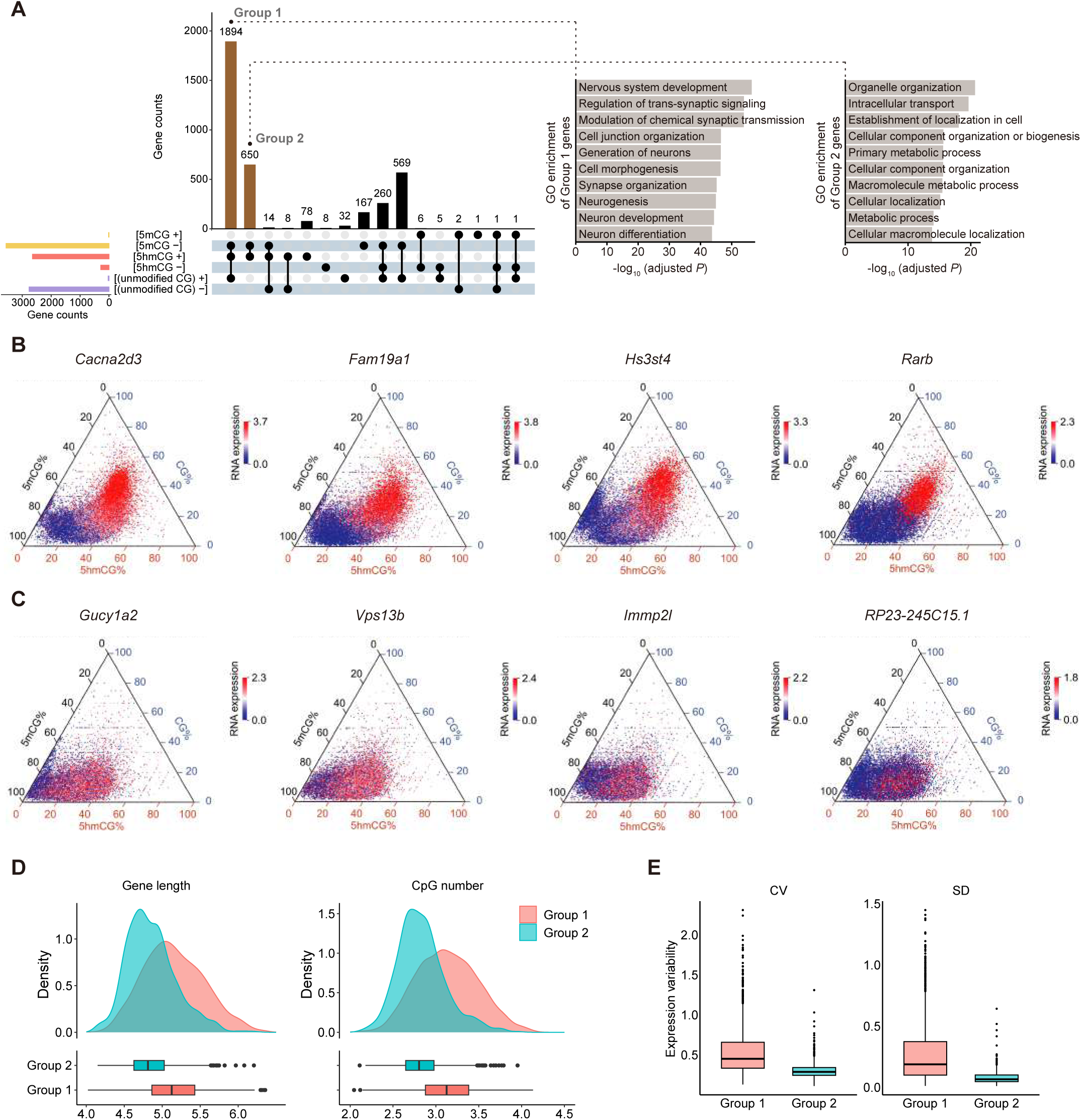
Diverse functions of DNA methylation and hydroxymethylation. (**A**) Left, upset plot showing different groups of genes with distinct gene regulatory relationships of various DNA modifications. Right, the Gene Ontology (GO) enrichment results of group 1 and group 2 genes. (**B, C**) Ternary plots showing the gene activating mechanism of group 1 genes and group 2 genes, related to **Fig. 2F**. (**D**) Distribution of the gene lengths and CpG numbers of genes in group 1 and in group 2. (**E**) Distribution of coefficients of variation (CVs) and standard deviations (SDs) of genes in group 1 and in group 2.

**Fig. S7.**
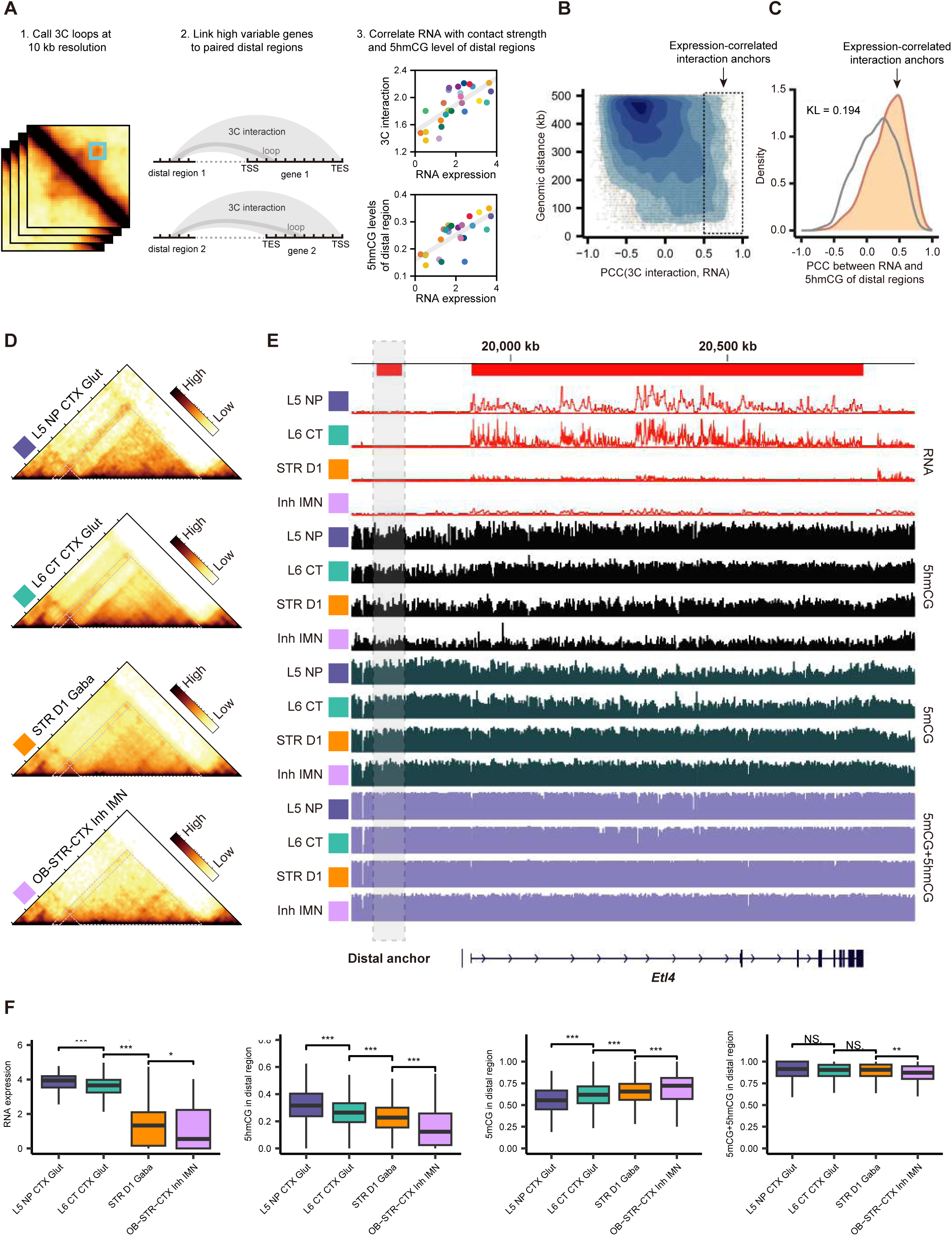
Distal regulation of DNA hydroxymethylation. (**A**) Workflow for analyzing DNA modifications in distal regulatory regions. (**B**) Identification of expression-correlated interaction anchors, where the Pearson correlation coefficients between contact strengths and RNA expression levels are required to be greater than 0.5. (**C**) Distribution of PCC values between RNA expression levels and 5hmCG levels at corresponding distal anchors. Expression-correlated interaction anchors (orange curve) show a more positive correlation compared to other interaction anchors (gray curve), with KL-divergence (bin size = 0.1) displayed in the figure. (**D**) Contact maps illustrating heterogeneous interaction strengths across different subclasses. (**E**) IGV snapshot showing *Etl4* RNA expression levels along with 5hmCG, 5mCG, and 5mCG+5hmCG levels of the distal regulatory region (chr2: 19,690,000–19,750,000). (**F**) Boxplots showing the *Etl4* RNA expression and DNA modification levels in the distal regulatory region.

**Fig. S8.**
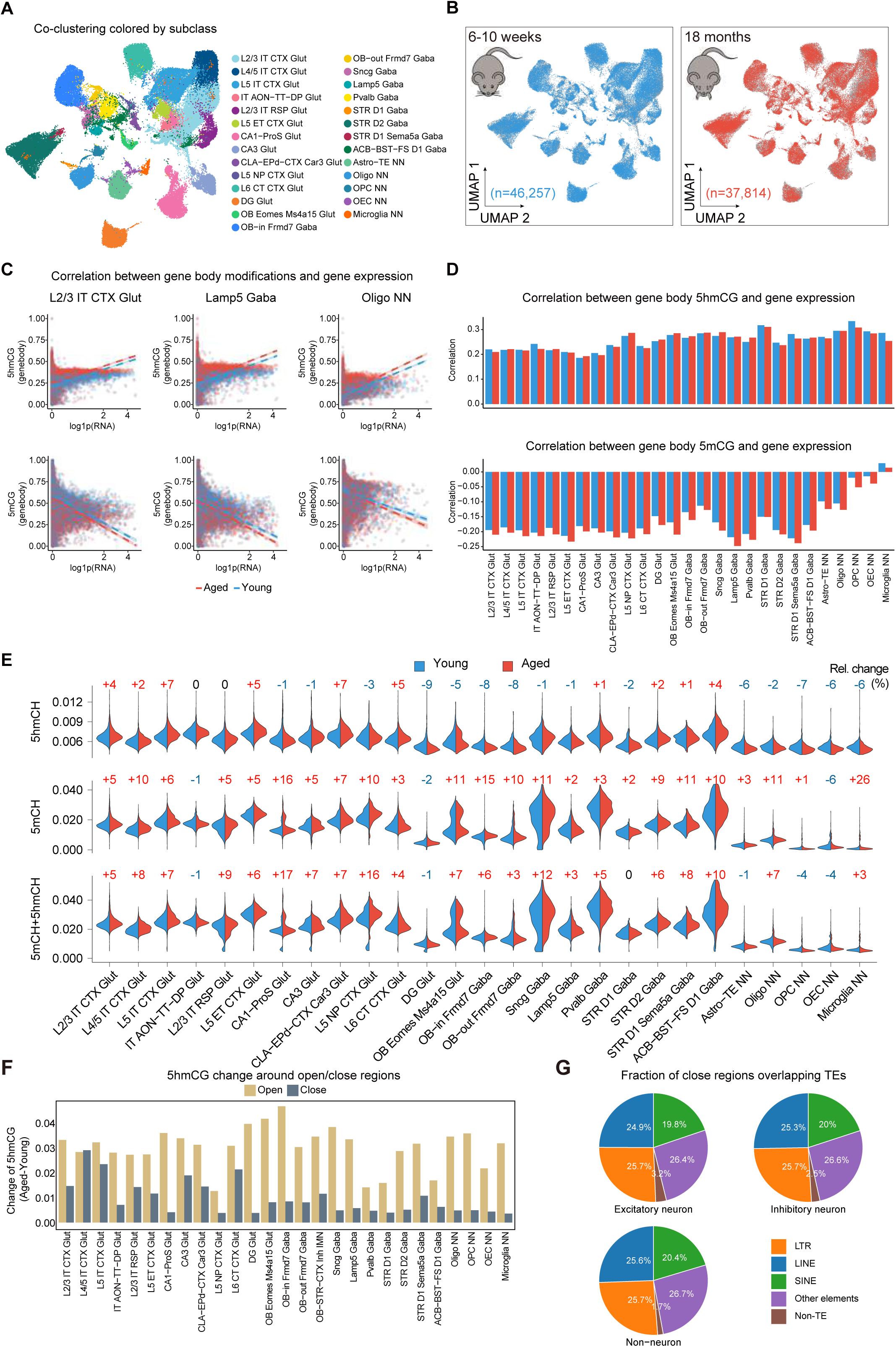
Aging-associated DNA modification changes. (**A-B**) UMAP visualization of transcriptome-based co-clustering of aged and young brain nuclei, colored by subclass labels (**A**) or mouse ages (**B**). (**C**) Scatter plots showing the relationships of 5hmCG and 5mCG levels with gene expression levels in genebodies of genes in different subclasses, in young (blue) and aged (red) mice. (**D**) PCCs between genebody 5hmCG and gene expression of genes in different subclasses in young (blue) and aged (red) mice, corresponding to **C**. (**E**) Global DNA modification ratios (5hmCH, 5mCH, and 5mCH+5hmCH) across subclasses demonstrate age-related alterations, with relative rates of change displayed at the top of each comparison, related to **Fig. 5B**. (**F**) Aging-related 5hmCG increments in open and closed chromatin regions across subclasses. (**G**) Fraction of close regions overlapping retrotransposable elements (TEs) in excitatory neuron, inhibitory neuron and non-neuron.

**Fig. S9.**
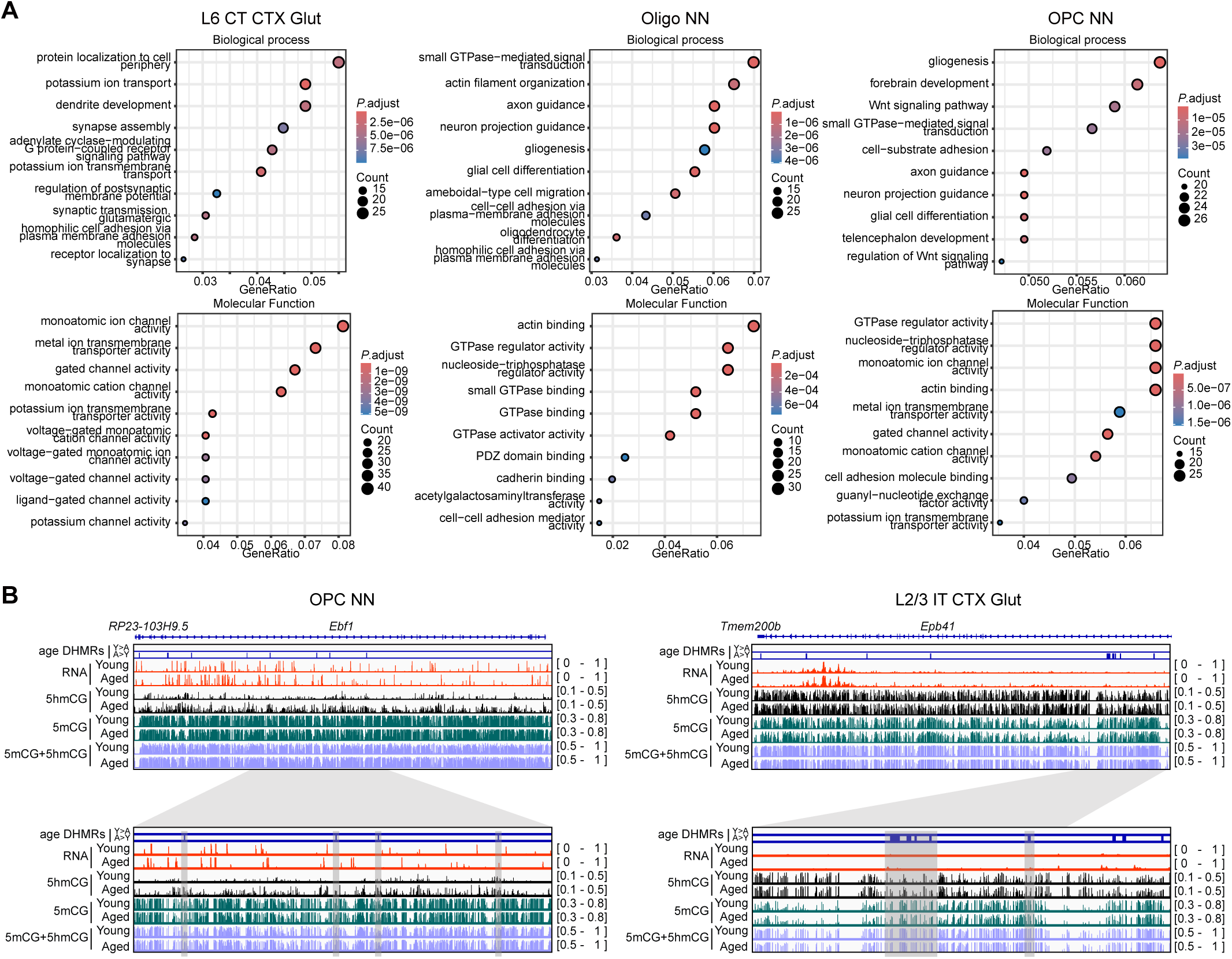
Aging-associated hyper-DHMRs reflecting subclass identities and aging-related gene expression changes. (**A**) GO analysis of the genes associated with top 1,000 aging-related hyper-DHMRs in ‘L6 IT CTX Glut’, ‘Oligo NN’, and ‘OPC NN’, showing enriched biological processes (BPs) and molecular functions (MFs). (**B**) IGV tracks showing two examples of coordinated 5hmCG accumulation and RNA upregulation during aging. The tracks in the bottom show zoom-in views with aging-related hyper-DHMRs highlighted in gray shadows.

## Supplementary Tables

Table S1. Primer sequences.

Table S2. Metadata of adult and aged mouse brain nuclei.

Table S3. Marker genes used to annotate Joint-Cabernet data.

Table S4. GO analysis of Group 1 and Group 2 genes.

Table S5. Cell-type-specific hyper-DHMRs overlapping gene bodies.

Table S6. Expression-associated interaction anchors with hyper-DHMRs.

Table S7. Location imputation of Joint-Cabernet profiles.

